# Dynamic S-acylation controls CMG2 maturation, extracellular matrix regulation, and anthrax toxin entry

**DOI:** 10.64898/2026.01.06.697925

**Authors:** Laurence Abrami, Sanja Blaskovic, Octave Joliot, Vincent Mercier, Béatrice Kunz, Francisco S. Mesquita, F. Gisou van der Goot

## Abstract

CMG2/ANTXR2 functions as a Collagen VI receptor and the primary portal for anthrax toxin entry. We find that CMG2 is regulated by ordered cycles of S-acylation and deacylation throughout its life cycle. Following synthesis in the endoplasmic reticulum, acylation by ZDHHC7 on two juxtamembrane cysteines protects folding intermediates from ER-associated degradation, resulting in a 5-fold increase in CMG2 biogenesis. The cytosolic thioesterase APT2 can remove these acyl chains, thereby controlling CMG2 levels. In the Golgi, CMG2 acylation by ZDHHC3 on a third cysteine to permit Arf6-dependent transport to the plasma membrane. At the cell surface, S-acylated CMG2 recruits APT2 in response to ligand binding, enabling release from the actin cytoskeleton and endocytosis. Accordingly, blocking APT2 suppresses the intracellular delivery of anthrax toxin, and inhibits CMG2-dependent Collagen VI degradation. These results define S-acylation–deacylation cycles as key regulators of CMG2 biogenesis and function, and highlight APT2 inhibition as a strategy to modulate CMG2 levels or prevent anthrax intoxication.

## INTRODUCTION

Capillary Morphogenesis Gene 2 (CMG2/ANTXR2) is a ubiquitously expressed type I membrane protein that plays a central role in extracellular matrix (ECM) homeostasis ^1–4^. Loss-of-function mutations in CMG2 cause Hyaline Fibromatosis Syndrome (HFS), a severe, often lethal, genetic disease characterized by excessive accumulation of collagen type VI (COL6) and other ECM components in various tissues, leading to organ dysfunction ^4–7^. CMG2 also serves as the major mammalian receptor for the anthrax toxins, providing entry of the enzymatic moieties of the toxin into the cell ^8^. Thus, a better understanding of the biogenesis and regulation of CMG2 is likely to provide key insights into the physiological role of this protein and its pathological role when hijacked by anthrax.

We previously reported that CMG2, and its paralog TEM8 (ANTXR1) also an anthrax toxin receptor ^9^ and also associated with a genetic disease^10^, undergo S-acylation and that this is required for productive anthrax toxin internalization ^11^. S-acylation refers to the covalent attachment of medium-chain fatty acids, typically palmitate, to cytosolic cysteine residues via thioester bonds ^12–14^. This process is catalysed by the transmembrane ZDHHC family acyltransferases and reversed by cytosolic acyl-protein thioesterases such as APT1/LYPLA1, APT2/LYPLA2, and ABHD17 family members ^13–15^. This dynamic modification can regulate protein conformation, subcellular trafficking, protein–protein interactions, turnover, and function ^16^. While we showed that acylation is involved in timing anthrax toxin endocytosis ^11^, the mechanistic consequence of S-acylation and its dynamics remain unknown, as do the enzymes involved.

CMG2 is synthesized with a signal sequence leading to its co-translational insertion into the membrane of the endoplasmic reticulum (ER)^17^. There, it undergoes folding, in particular through the help of glycan-binding chaperones ^17^. However, the CMG2 ectodomain folds inefficiently, rendering the protein vulnerable to genetic mutations ^18^. A variety of Hyaline Fibromatosis Syndrome missense mutations impair CMG2 folding, resulting in its degradation via the ER-associated degradation (ERAD) and protein-loss dependent loss-of-function ^3,18,19^. Little is however known about the folding and maturation of CMG2 in the ER.

Once folded, CMG2 exits the ER and transits through the Golgi to reach the plasma membrane ^18^. At the cell surface, ligand-free CMG2 associates with the cytoskeleton by interacting with a Talin-Vinculin-Actin (TVA) containing complex ^20^. This TVA interaction is released when a ligand, such anthrax toxin or Collagen VI ^4^ binds to the ectodomain of CMG2, so that its cytoplasmic domain can then recruit proteins recruited for CMG2 signaling and endocytosis such as RhoA, β-arrestin, and Cbl ^11,20,21^. This signalling transition is essential for both anthrax toxin uptake and CMG2-mediated internalization and lysosomal degradation of COL6 ^20^. A role of S-acylation in this transition has not been demonstrated.

Here we reveal that various stages of the CMG2 life cycle, including maturation in the ER, trafficking and signalling, are all governed by sequential S-acylation-deacylation cycles. We find that newly synthesized CMG2 is protected from ERAD by ZDHHC7-mediated S-acylation of juxtamembrane cysteines, which increases the flux of folded proteins exiting the ER. This S-acylation is however counteracted in the ER by APT2. Its inhibition indeed enhances CMG2 expression in cultured cells and in mice, identifying APT2 as a promising therapeutic target for HFS alleles that affect CMG2 folding. After ER exit, upon reaching the Golgi, ZDHHC3-dependent S-acylation of a distal cysteine enables Arf6-dependent CMG2 trafficking to the plasma membrane. At the cell surface, only fully S-acylated CMG2 is capable of propagating the ligand-induced conformational change across the membrane to the cytosolic tail. Ligand binding in particular triggers recruitment of APT2 and CMG2 deacylation, enabling release from the TVA complex and recruitment of RhoA and β-arrestin, converting the receptor into an internalization-competent state. As a consequence, genetic or chemical inhibition of APT2 blocks anthrax toxin endocytosis and subsequent cellular damage.

These findings identify sequential S-acylation steps that govern all steps of the CMG2 life cycle from maturation, to trafficking and ligand-induced signalling, and establish APT2 as a key druggable regulator of CMG2 function in both HFS and anthrax intoxication.

## RESULTS

### CMG2 undergoes sequential S-acylation by ZDHHC7 and ZDHHC3

It has been reported that CMG2 and its paralogue TEM8 undergo S-acylation (Abrami et al., 2006, Fig. 1a). We confirm here that CMG2 indeed undergoes S-acylation in cultured cells using [³H]-palmitate metabolic labelling, and in various mouse tissues using the Acyl-RAC assay which captures S-acylated proteins in protein extracts (Fig. 1bc). CMG2 isoform 4 contains three cytosolic cysteines (Cys344, Cys345, and Cys379; Fig. 1a). To identify which residues are modified, we generated a panel of single, double, and triple cysteine-to-alanine mutants, with the wild-type (WT) corresponding to CCC and the triple C344A-C345A-C379A mutant denoted as AAA. Loss of all three cysteines abolished [³H]-palmitate incorporation (Fig. 1d). Among the double mutants, the AAC variant (where Cys344 and Cys345 are mutated) showed the strongest reduction, indicating that acylation of these two juxtamembrane cysteines influences acylation of the third site. Kinetic labelling experiments confirmed that acylation is initiated at Cys344/345 and subsequently proceeds to Cys379 (Fig. 1e).

**Figure 1:**
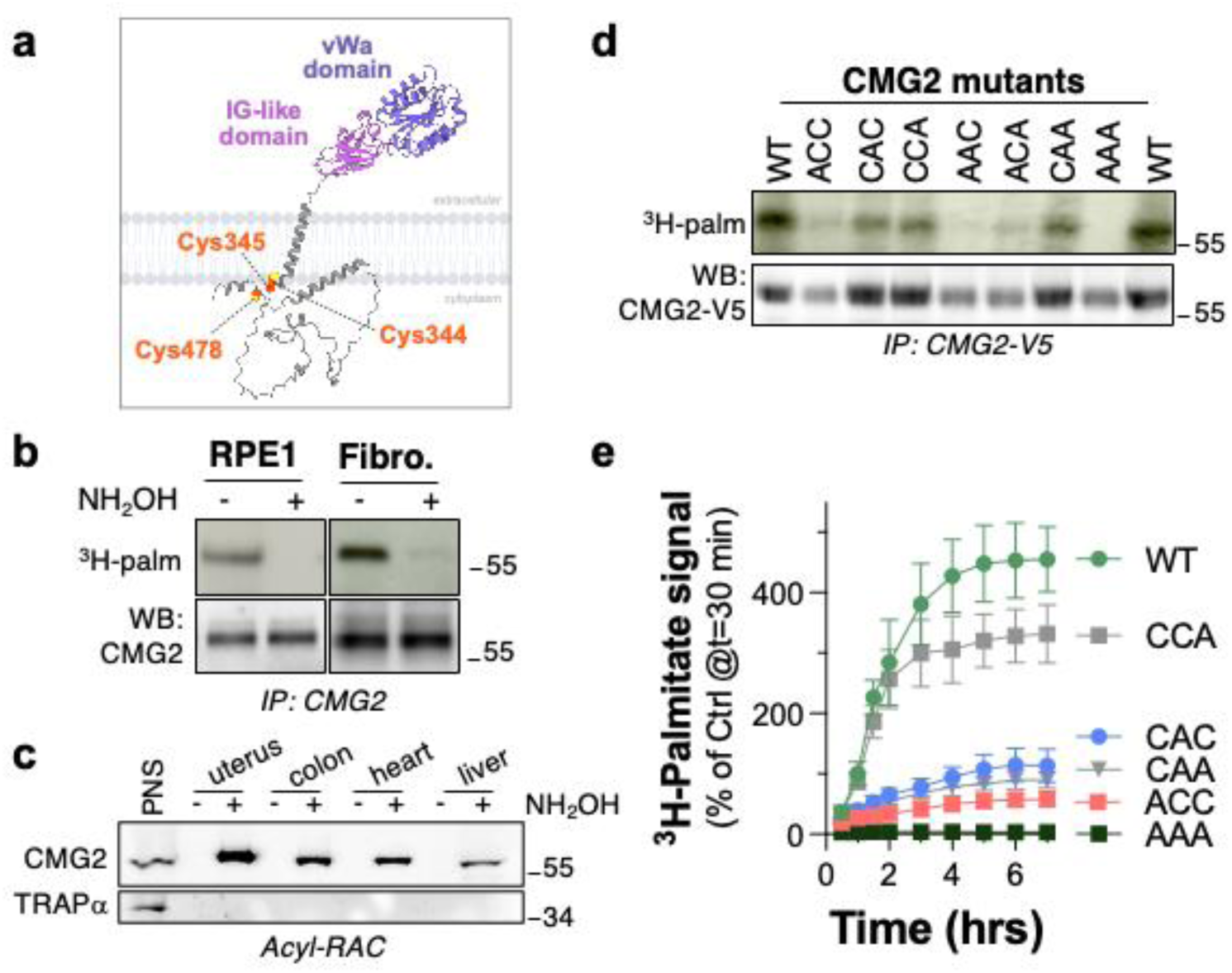
CMG2 is S-palmitoylated on Cys-344, Cys-345 and Cys-479. **a**. Schematic representation of CMG2 showing the von Willebrand A domain, Ig-like domain, transmembrane region, and cytosolic tail. The three cytosolic cysteines are indicated. **b**. RPE1 cells or primary human fibroblasts were metabolically labeled for 2 h at 37 °C with [³H]-palmitic acid. CMG2 was immunoprecipitated and samples were treated or not with hydroxylamine (+NH₂OH) to cleave thioester-linked acyl chains. Proteins were separated by SDS–PAGE and analyzed by autoradiography (³H-palm) and by immunoblotting against CMG2. **c**. CMG2 in mouse organs was analyzed using Acyl-RAC. Equal volumes of input (Inp.) and Acyl-RAC eluates (+NH₂OH) were separated by 4–20% SDS–PAGE. Immunoblotting was performed with anti-mouse CMG2 (monoclonal 8F7) or anti-TRAPα as a control.**d**. HeLa cells were transfected for 24 h with WT and mutants CMG2-V5. Cells were labeled for 2 h with [³H]-palmitate at 37 °C and CMG2-V5 was immunoprecipitated. Autoradiography and anti-V5 immunoblotting were used to detect acylated and total CMG2, respectively. **d.** HeLa cells expressing WT or cysteine mutants were labeled for different times with [³H]-palmitate at 37 °C. CMG2-V5 was immunoprecipitated and analyzed as in (d). ³H-palmitate incorporation at 1 h was set to 100%, and each timepoint expressed relative to this value (n=3, mean ± SD).

To identify the acyltransferases responsible for CMG2 modification, we performed an siRNA screen targeting all 23 human ZDHHC enzymes, grouped into six pools. Only pool M1 led to a reduction in CMG2 acylation (Fig. 2a, Extended Data Fig. 1a). Subsequent individual knockdowns within this pool revealed that ZDHHC7 and ZDHHC3 are responsible for mediating CMG2 S-acylation (Fig. 2b, Extended Data Fig. 1b). These two enzymes also S-acylate the 4 cysteines present in TEM8 ^11^ (Extended Data Fig. 1c).

**Figure 2:**
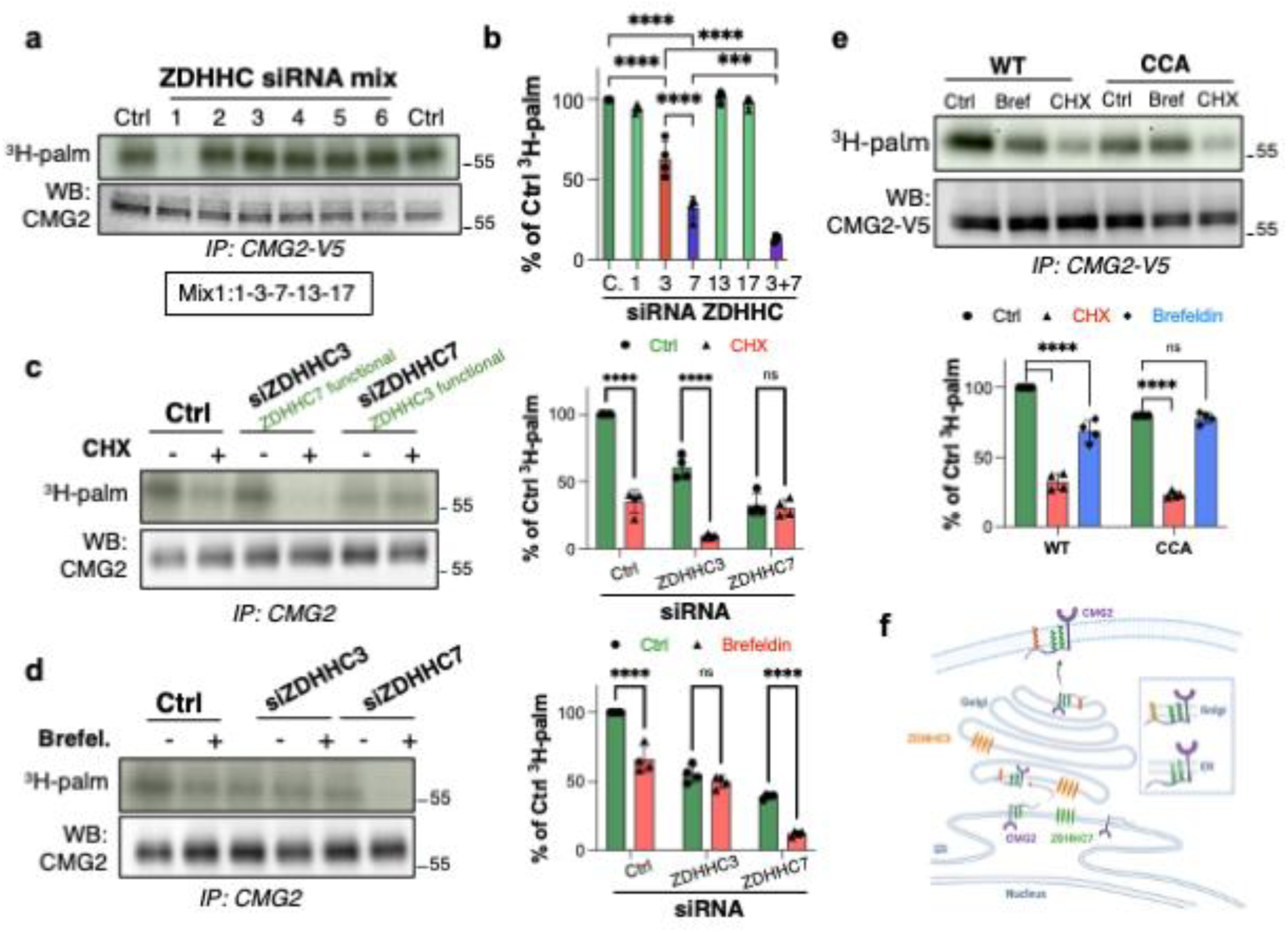
CMG2 S-palmitoylation is mediated by ZDHHC3 and ZDHHC7. **a,b.** RPE1 cells were silenced with pooled (a, the composition of the pools is provided in the Methods section) or individual (b) siRNAs targeting human ZDHHC enzymes, or non-targeting control siRNA. Cells were metabolically labeled for 2 h at 37 °C with [³H]-palmitic acid, CMG2 was immunoprecipitated, and samples were analyzed by SDS–PAGE followed by autoradiography and anti-CMG2 immunoblotting. [³H]-palmitate incorporation was quantified by Typhoon imaging and normalized to control = 100% (a: n=3; b: n=4; mean ± SD). **c-e**. RPE1 cells were silenced with siZDHHC3, siZDHHC7, or control siRNA. For panel (**e**), HeLa cells were transfected with WT or CCA CMG2-V5. Cells were pre-treated or not with cycloheximide (CHX) (**c,e**) or Brefeldin A (**d,e**) for 1 h and maintained under the same conditions during metabolic labeling. Cells were labeled for 2 h at 37 °C with [³H]-palmitate, CMG2 was immunoprecipitated with anti-CMG2 (c,d) or anti-V5 (**e**), and samples were analyzed by SDS–PAGE, autoradiography (³H-palm), and immunoblotting with anti-CMG2 (**c,d**) or anti-V5 (**e**). Radiolabel incorporation was quantified by Typhoon imaging and normalized to control siRNA/untreated = 100% (n=4; mean ± SD). **f.** Sequential CMG2 S-acylation occurs first on Cys344/345 by ZDHHC7 in the ER, followed by Cys379 modification by ZDHHC3 in the Golgi.

**Extended Data Figure 1:**
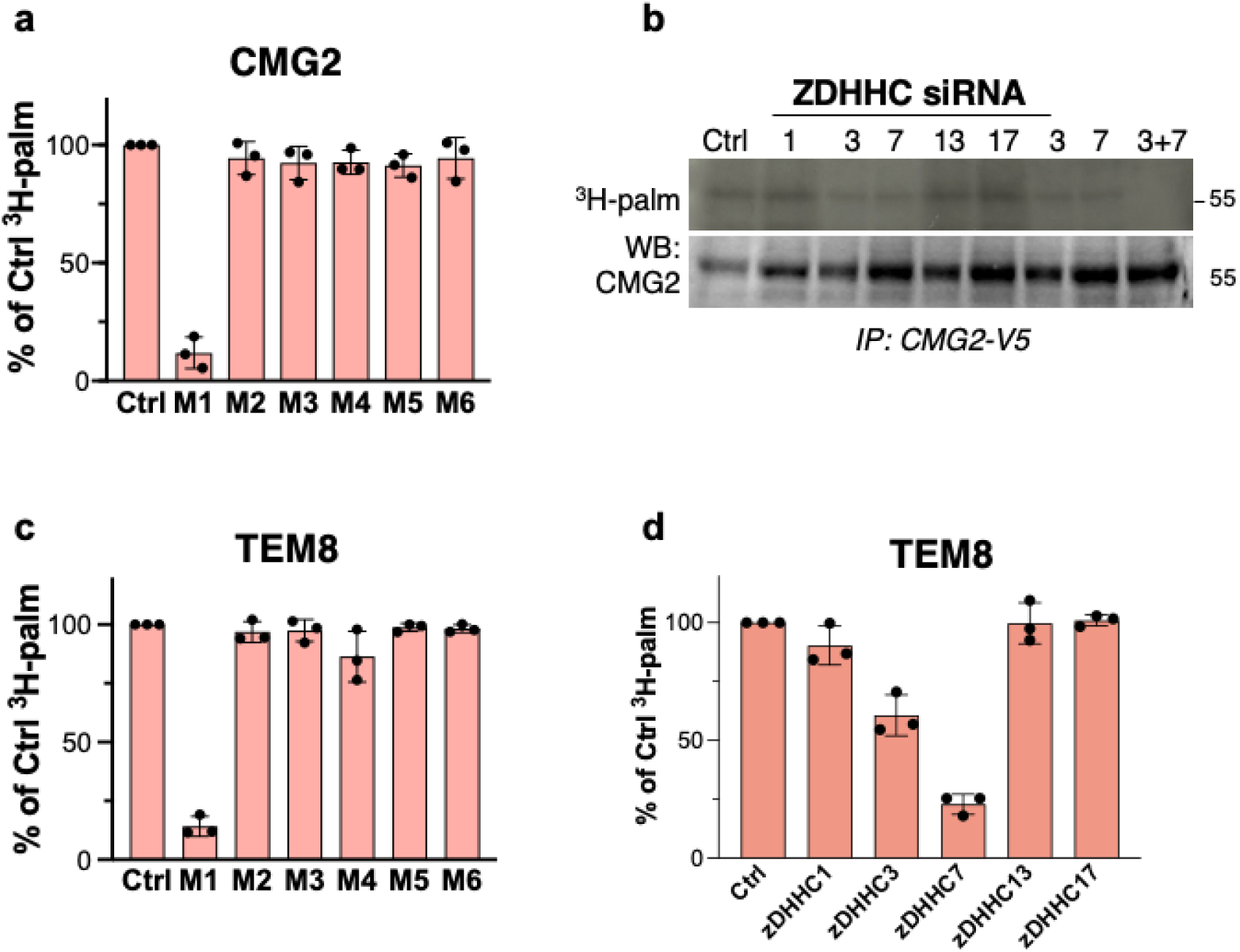
ZDHHC3/7 are the primary acyltransferases for CMG2 and TEM8. **a,b.** RPE1 cells were silenced for 3 days with pooled (a) or individual (b) siRNAs targeting human ZDHHC enzymes, or control siRNA. Cells were metabolically labeled for 2 h at 37 °C with [³H]-palmitic acid, CMG2 was immunoprecipitated with goat anti-human CMG2, and samples were analyzed by SDS–PAGE followed by autoradiography (³H-palm) and anti-CMG2 immunoblotting. [³H]-palmitate incorporation was quantified with a Typhoon scanner and normalized to control = 100% (n=3, mean ± SD). **c.** HeLa cells were silenced for 3 days with pooled or individual siRNAs against human ZDHHC enzymes, metabolically labeled with [³H]-palmitate for 2 h at 37 °C, and TEM8 was immunoprecipitated with goat anti-TEM8. Samples were analyzed by SDS–PAGE, autoradiography, and anti-TEM8 immunoblotting. Radiolabel incorporation was quantified by Typhoon imaging and normalized to control = 100% (n=3, mean ± SD).**d.** Dual silencing of ZDHHC3 and ZDHHC7 markedly reduced [³H]-palmitate incorporation into CMG2 and TEM8, confirming that these two acyltransferases are the principal determinants of receptor palmitoylation. Quantification was performed as in (a-c).

We next investigated whether the two enzymes act on different cysteines, and possibly in different intracellular compartments. ZDHHC7 localizes to the early secretory pathway (ER and Golgi) and ZDHHC3 to the Golgi ^22^. To investigate whether acylation occurs in the ER when CMG2 is newly synthesized, we inhibited protein synthesis with cycloheximide. We monitored ZDHHC7-mediated S-acylation by silencing ZDHHC3, and conversely ZDHHC3-mediated S-acylation by silencing ZDHHC7. ZDHHC7-mediated acylation was essentially abolished by cycloheximide, whereas ZDHHC3-mediated acylation was unaffected (Fig. 2c), consistent with its Golgi localization. Next, we tested the effect of disrupting the Golgi with brefeldin A, and observed the opposite pattern: ZDHHC7-mediated S-acylation was unaffected by brefeldin A, whereas ZDHHC3-mediated S-acylation was essentially abolished (Fig. 2d). Finally, we tested whether the CCA (i.e. C379A only) mutation would recapitulate ZDHHC3 silencing. Cycloheximide treatment strongly reduced CCA S-acylation while brefeldin A treatment had no effect (Fig. 2e). Thus, CMG2 undergoes sequential S-acylation, first in the ER on Cys344 and Cys345 by ZDHHC7, and subsequently on Cys379 by ZDHHC3 in the Golgi (Fig. 2f).

### S-acylation-deacylation controls maturation of CMG2 in the ER

Because CMG2 is S-acylated during or shortly after synthesis, we asked whether this modification influences its biogenesis and thus the amount of fully folded protein that exits the ER. We performed [³⁵S]-Cys/Met pulse-chase labelling with a short 20 min pulse to focus on newly synthesized molecules. CMG2 decayed with a t₁/₂ of ∼5 h, both for endogenous (Ctrl) and ectopically expressed (WT) protein (Fig. 3a). Inhibition of the proteasome with MG132 increased the signal at t=0 and markedly delayed and reduced degradation, indicating that in untreated cells, a major fraction of newly synthesized CMG2 is destroyed during the first 5-6hrs (maturation phase, Fig. 3a) by ERAD, reminiscent of the behaviour of the ΔF508 cystic fibrosis transmembrane regulator CFTR mutant ^23,24^. When we performed a similar experiment on the AAA triple mutant, degradation of the newly synthesized protein was significantly accelerated (t₁/₂ ≈2 h) and more pronounced, with degradation already significant during the pulse period (Fig. 3a). Silencing ZDHHC7 and 3 completely recapitulated the effect of mutating the three cysteines (Extended Data Fig. 2a). Accelerated degradation following the metabolic pulse was also observed when mutating only Cys344 or Cys345 (Fig. 3b, normalised to t=0 to facilitate kinetic comparison), consistent with these residues being S-acylated in the ER and protecting folding intermediates from ERAD.

**Figure 3:**
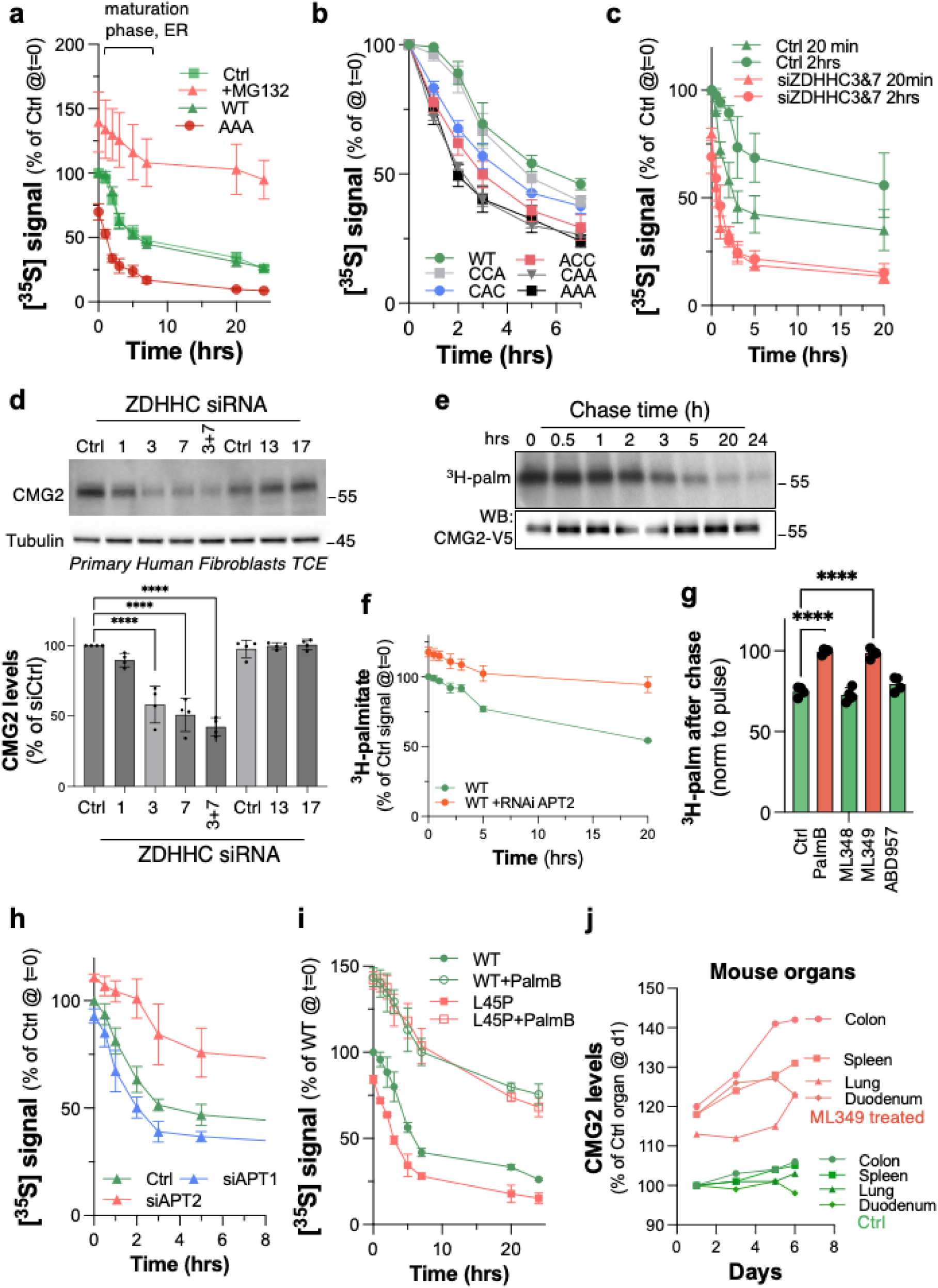
ZDHHC3 and ZDHHC7 control CMG2 biogenesis. **a-c**. HeLa cells expressing WT or mutant CMG2 (**a,b**), or RPE1 cells silenced with individual ZDHHC siRNAs (**c**), were pulse-labeled for 20 min with [³⁵S]-Met ± MG132 (**a**) and chased for the indicated times. CMG2 was immunoprecipitated and analyzed by SDS–PAGE, autoradiography and anti-CMG2 immunoblotting. Radiolabel incorporation was quantified (Typhoon) and normalized to untreated WT/control = 100% (n=3, mean ± SD). **d**. Primary fibroblasts silenced for 72 h with the indicated siRNAs were analyzed by immunoblot of total cell extracts (40 µg) for endogenous CMG2 ± Tubulin loading control (n=4, mean ± SEM). **e-g**. HeLa cells expressing CMG2-V5 (e) or RPE1 cells treated with Palmostatin B, ML348, ML349 or ABD957 (g) were labeled 2 h with [³H]-palmitate and chased as indicated. For APT2 involvement, RPE1 cells were silenced with siAPT2 (f). CMG2 was immunoprecipitated and analyzed by autoradiography and immunoblot. [³H]-palmitate signal after chase was normalized to pulse timepoint (n=3, mean ± SD). **h, i**. Effects of deacylation inhibition on folding. RPE1 cells silenced for APT1/2 (h) or HeLa expressing WT or L45P CMG2 ± Palmostatin B (i) were pulse-labeled 20 min with [³⁵S]-Met and chased. CMG2 was immunoprecipitated, quantified, and expressed relative to control = 100% (n=3, mean ± SD). **j**. In vivo stabilization. Mice were treated i.p. with ML349 (40 mg/kg; days 0,2,4,6,8), organs collected at indicated timepoints, and endogenous CMG2 measured by immunoblot of tissue lysates (40 µg).

**Extended Data Figure 2:**
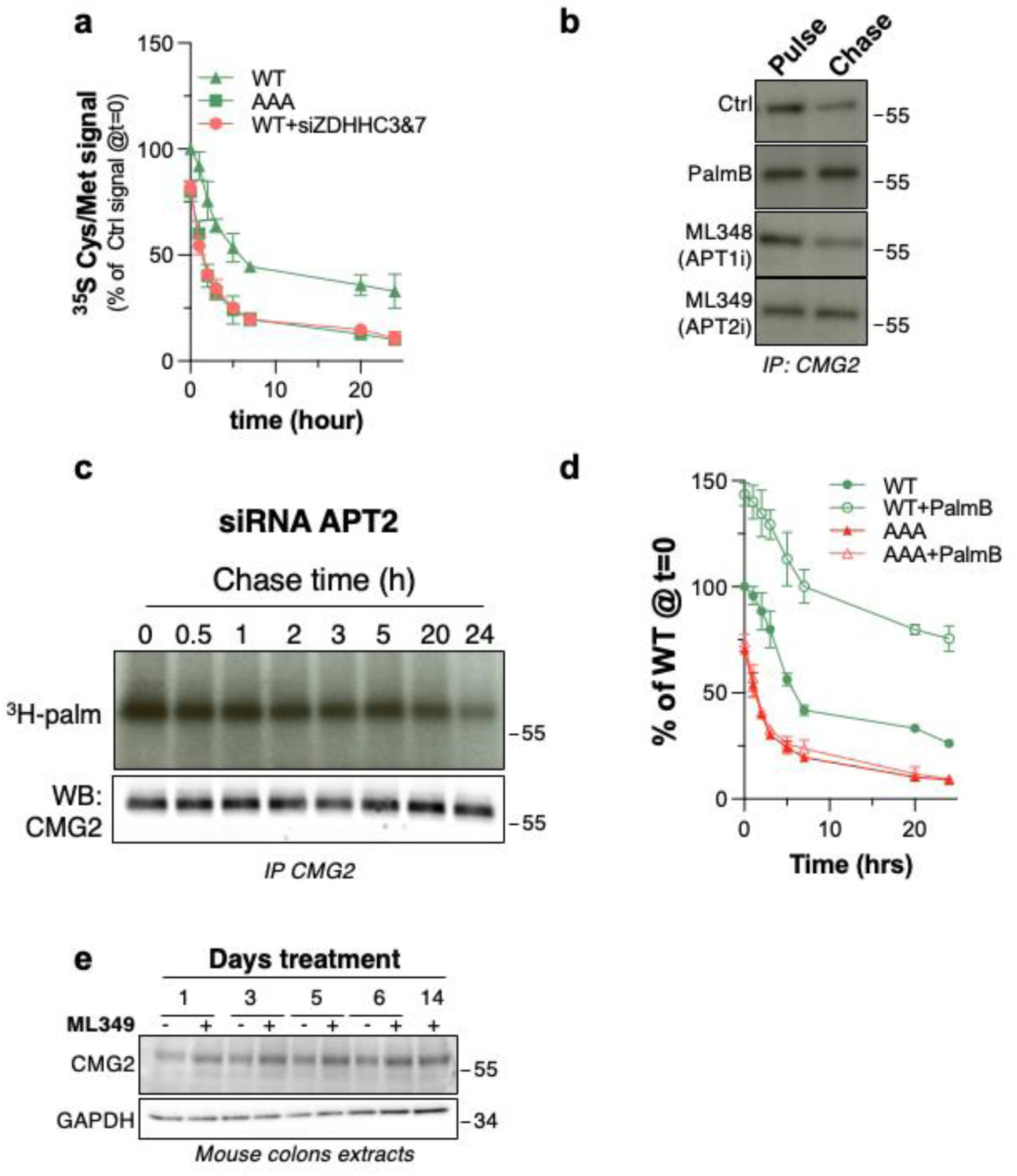
S-Acylation–deacylation of CMG2. **a.** HeLa cells expressing CMG2-V5 WT or AAA were silenced for 72 h with siZDHHC3+siZDHHC7 or control RNAi, pulse-labeled 20 min with [³⁵S]-Met and chased as indicated; CMG2-V5 was immunoprecipitated and analyzed by SDS–PAGE, autoradiography and immunoblotting, and radiolabel incorporation quantified (Typhoon) and normalized to WT/no RNAi = 100% (n=3, mean ± SD). **b.** RPE1 cells were treated 4 h before and during labeling with Palmostatin B (50 µM), ML348 (10 µM) or ML349 (10 µM), labeled 3 h with [³H]-palmitate and chased 5 h; CMG2 immunoprecipitates were analyzed by SDS–PAGE and autoradiography. **c**. RPE1 cells were transfected with siAPT2 for 72 h, labeled 2 h with [³H]-palmitate and chased; CMG2 was immunoprecipitated and analyzed by SDS–PAGE, autoradiography and immunoblotting. **d.** HeLa cells expressing CMG2-V5 WT or AAA were treated ± Palmostatin B (50 µM; 4 h before and during labeling), pulse-labeled 20 min with [³⁵S]-Met and chased; CMG2-V5 decay was quantified as in a. and normalized to WT/no drug = 100% (n=3, mean ± SD). **e.** Mice received ML349 intraperitoneally (40 mg/kg on days 0, 2, 4, 6, 8); colon was collected at indicated times and CMG2 abundance assessed by immunoblotting of total lysates (40 µg) with anti-CMG2 and anti-GAPDH.

The difference in the [³⁵S]-Cys/Met signal at t=0 for WT and AAA CMG2 indicates that some WT molecules undergo S-acylation during the 20 min pulse. However S-acylation of the pool of newly synthesized CMG2 molecules may take longer than 20 min. To test this, we compared the effects of 20 min and 2 h [³⁵S]-Cys/Met labeling durations. Extending the pulse period resulted in reduced and delayed degradation of the pool of synthesized WT CMG2 (Fig. 3c). The effect of the pulse length was lost when silencing ZDHHC7 and ZDHHC3 (Fig. 3c), confirming that the stabilizing effect was due to S-acylation. We have made similar observations for two other S-transmembrane proteins undergoing S-acylation in the ER, calnexin and LRP6, and

### Deacylation by APT2 in the ER targets CMG2 to premature ERAD

Since S-acylation is reversible, we next examined whether during biogenesis in the ER, CMG2 also undergoes deacylation. To address this, we first assessed whether CMG2 can be deacylated at all. [³H]-palmitate pulse-chase analysis, using a 2 hrs pulse, showed a slow and progressive loss of radiolabel from CMG2 (Fig. 3ef). Loss of [³H]-palmitate signal can occur by deacylation as well as by protein degradation. To determine the contribution of deacylation, we tested the effects of acyl-protein thioesterase inhibitors. The pan-thioesterase inhibitor Palmostatin B and the APT2-specific inhibitor ML349, but not the APT1 inhibitor ML348 or the ABHD17 inhibitor ABD957, prevented the loss of [³H]-palmitate during the chase period (Extended Data Fig. 2b, Fig. 3g). Kinetic measurements showed that APT2 silencing indeed substantially delayed and reduced the loss of [³H]-palmitate from CMG2 (Fig. 3f, Extended Data Fig. 2c).

To more specifically determine whether APT2, which is a cytosolically expressed enzyme, can act on CMG2 during biogenesis, we again performed [³⁵S]-Cys/Met pulse-chase experiments. Knockdown of APT2 protected newly synthesized CMG2 from ERAD, with stabilization evident already during the pulse (Fig. 3f). The effect was even more striking when cells were treated with Palmostatin B, for reasons that remain to be clarified (Extended Data Fig. 2d). Importantly, Palmostatin B did not affect the decay kinetics of the AAA mutant, indicating that the observed effects are related to S-acylation of WT CMG2 (Extended Data Fig. 2d).

These results show that during the folding of a newly synthesis population of CMG2, APT2 can remove the protective S-acylations, thereby promoting the premature targeting of these molecules to ERAD. Quantitatively, comparison of CMG2 levels 5 h after the pulse (Fig. 3c,h) reveal an approximately five-fold difference between ZDHHC7/3 silencing and APT2 silencing, highlighting the major contribution of S-acylation to CMG2 folding efficiency.

This mechanism has implications for Hyaline Fibromatosis Syndrome (HFS) missense mutations that affect folding ^3,18,19^. We previously showed that several of these mutations, which map to the extracellular von Willebrand A domain, fold properly when produced in *E. coli* ^18^, indicating that in human cells it is the slower folding kinetics – rather than an intrinsic inability to fold–that triggers ERAD recognition. Our current findings suggest that inhibiting deacylation, to protect folding intermediates from ERAD targeting, should lead to an increase in the amount of successfully folded receptors. We tested this hypothesis on the L45P HFS mutation ^18^. Indeed, Palmostatin B prevented the premature degradation of CMG2 L45P and led to an increase of CMG2 levels, reaching similar levels as observed for WT CMG2 (Fig. 3i). As a further proof of concept for this therapeutic avenue to increase the CMG2 protein levels, we treated mice by intraperitoneal injection with the APT2 inhibitor ML349 for several days, sacrificed the animals at defined time points, and collected multiple organs for analysis. ML349 treatment led to a time-dependent increase in CMG2 abundance across all tissues tested, as most apparent in the colon (Fig. 3j, Extended Data Fig. 2e). Altogether, the present findings show that newly synthesized CMG2 is robustly protected from ERAD through the S-acylation of juxtamembrane cysteines and indicate that inhibition of APT2 could be a potential strategy for improving CMG2 expression in contexts where receptor folding is impaired.

### CMG2 trafficking from Golgi to the plasma membrane is ZDHHC3-dependent and Arf6-mediated

We next assessed what the functional role of ZDHHC3-mediated S-acylation of CMG2 could be. In control cells, CMG2 was predominantly detected at the plasma membrane, whereas ZDHHC3 knockdown caused a marked accumulation of CMG2 in the Golgi (Fig. 4a), a result confirmed by high-content image analysis (Extended Data Fig. 3). Consistently, surface biotinylation revealed a strong reduction in CMG2 plasma membrane levels upon ZDHHC3 silencing (Fig. 4bc). To exclude enhanced endocytosis in ZDHHC3 silenced cells as an alternative explanation, we repeated the surface biotinylation assay in clathrin heavy chain (CHC)-silenced cells, to prevent CMG2 endocytosis ^21^. Under these conditions, the effect of ZDHHC3 depletion on CMG2 surface abundance was even more striking (Fig. 4c). Thus, the decrease in surface CMG2 upon ZDHHC3 silencing is due to a defect in anterograde trafficking to the plasma membrane rather than increased internalization.

**Figure 4:**
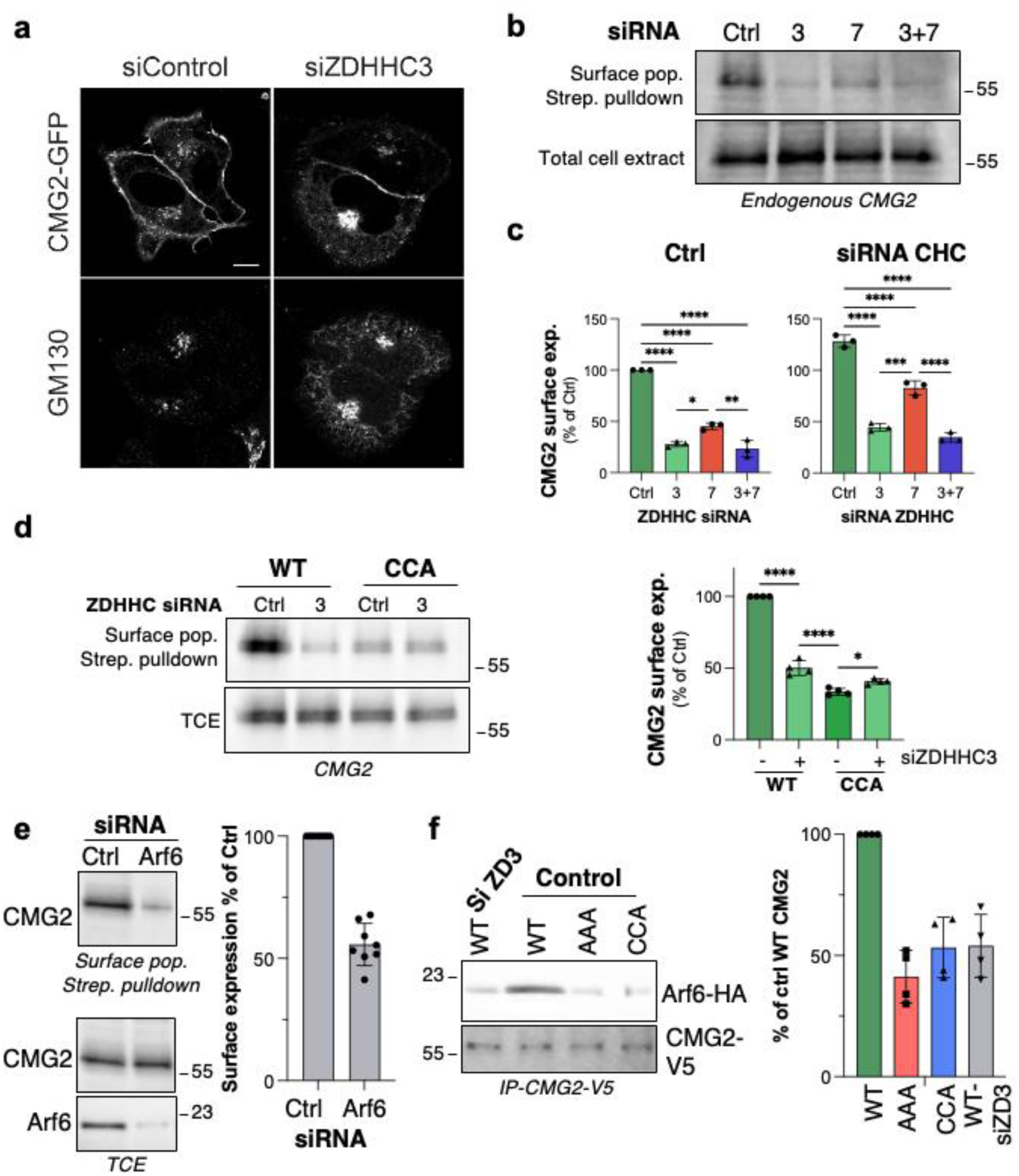
ZDHHC3-dependent, Arf6-mediated trafficking of CMG2 to the plasma membrane. **a.** Immunofluorescence microscopy of CMG2-GFP–expressing HeLa cells treated for 72 h with control or ZDHHC3 siRNA. GM130 marks the Golgi. Scale bar, 10 µm. **b.** RPE1 cells were silenced for 3 d with siZDHHC3, siZDHHC7, both, or control RNAi. Plasma membrane proteins were biotinylated, total lysates collected, and streptavidin pull-downs probed for endogenous CMG2 by western blot (n=3, mean ± SE). **c.** Quantification of biotinylated CMG2 surface levels under conditions in (b). **d.** HeLa cells were silenced for 3 d with siZDHHC3 or control RNAi, then transfected for 24 h with CMG2-V5 WT or CCA. Surface proteins were biotinylated and streptavidin precipitates probed with anti-V5 (n=4, mean ± SE). **e.** RPE1 cells were silenced for 3 d with siARF6 or control RNAi, plasma membrane proteins biotinylated, precipitated with streptavidin, and membranes blotted for endogenous CMG2 and ARF6 (n=8, mean ± SE). **f.** RPE1 cells were silenced for 3 d with siZDHHC3 or control RNAi, transfected 24 h with CMG2-V5 WT, CCA, or AAA together with ARF6-HA, immunoprecipitated using V5 beads, and blotted for CMG2-V5 and ARF6-HA (n=4, mean ± SE).

**Extended Data Fig. 3:**
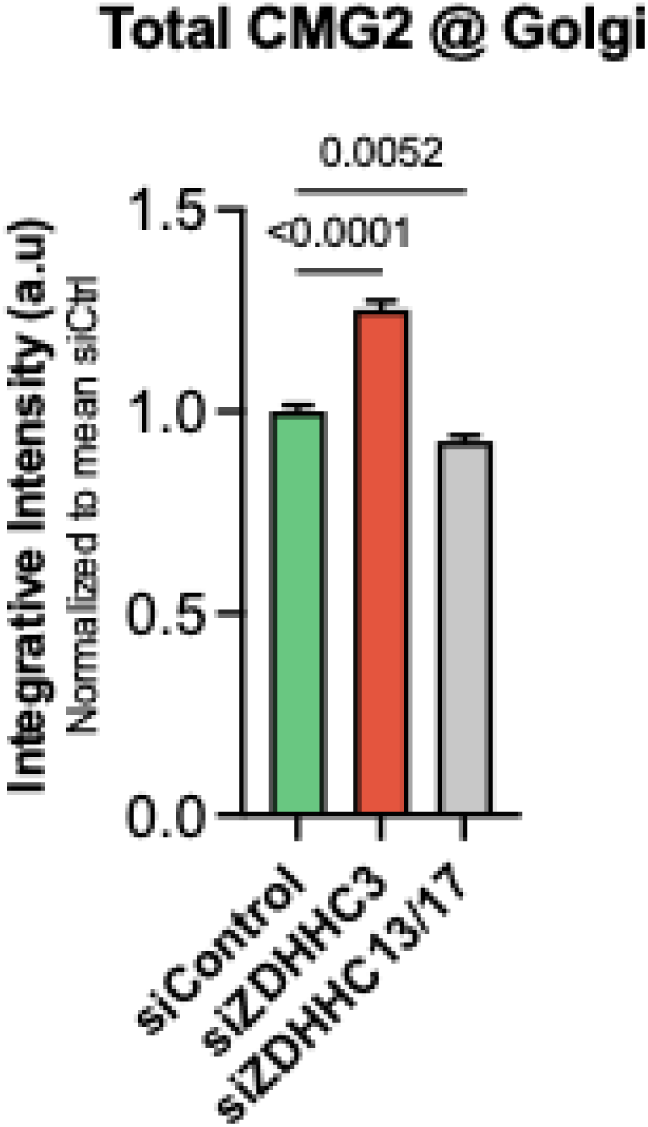
ZDHHC3 promotes exit of CMG2 from the Golgi. High-throughput automated immunofluorescence quantification of CMG2–GFP localization in HeLa cells transfected with control siRNA, siZDHHC3, or siZDHHC13/17 (72 h) and expressing CMG2–GFP for 24 h. Data represent integrative intensity values at the Golgi, defined by GM130-positive regions, and normalized to siCtrl. Bar graphs show mean values ± s.e.m. from one of two independent experiments with equivalent results. *n* = 2548 (siCtrl), 2540 (siZDHHC3), and 2756 (siZDHHC13/17) cells analysed. *p* values calculated by one-way ANOVA followed by Dunnett’s multiple-comparisons test against siCtrl.

Because ZDHHC3 S-acylates Cys379, we next analysed the CCA mutant. This mutant exhibited reduced plasma membrane localization and was insensitive to ZDHHC3 knockdown (Fig. 4d), confirming that S-acylation of Cys379 promotes CMG2 surface delivery. Since Golgi-to-plasma membrane transport of S-acylated proteins has been proposed to depend on Arf6 ^25^, we tested its potential role. Silencing Arf6 led to decreased CMG2 surface levels, and the interaction between CMG2 and Arf6 was abolished when Cys379 was mutated, or ZDHHC3 was depleted (Fig. 4e-f). Together, these findings show that ZDHHC3-mediated acylation of Cys379 allows CMG2 to engage Arf6 and traffic efficiently from the Golgi to the plasma membrane.

### S-acylated CMG2 can undergo ligand-induced cytoskeletal disengagement and signalling

We next investigated whether the S-acylation status of CMG2 influences its functions, first focusing on ligand-induced signalling at the plasma membrane ^20^. In the absence of ligand, surface CMG2 interacts with Talin and thereby Vinculin and Actin (TVA), anchoring it to the actin cytoskeleton ^20^. Ligand binding however triggers a switch from TVA binding to recruitment of the actin regulators RhoA and mDia, as well as with signalling and endocytic proteins such as the scaffold protein β-arrestin or src which then phosphorylates CMG2 ^20,26,27^ (Fig. 5a). This switch is readily visible when analysing CMG2 immunoprecipitates under conditions where cells are or not treated with the receptor binding component of the anthrax toxins, the protective antigen (PA), either in its 83 or 63 kDa form at 4°C followed by a 30 min incubation at 37°C: in control cells, Talin and actin co-IP with CMG2, while in toxin treated cells, the receptor is associated with RhoA (Fig. 5b). The TVA release is not observed when cells are kept at 4° ^20^. The AAA CMG2 mutant failed to release TVA components in response to PA binding, and consistently did not recruit RhoA (Fig. 5bc). Note that the presence of PA in the CMG2 IP indicates that transient transfection of AAA leads to sufficient expression of the mutant at the cell surface to monitor the binding. The CCA mutant exhibited a partial phenotype, with experiment-to-experiment variability, while the AAC mutant retained significant TVA binding and consistently showed limited RhoA recruitment (Fig. 5bc). Similarly, silencing ZDHHC7 and ZDHHC3 prevented PA-induced TVA release, despite PA binding, and RhoA recruitment (Fig. 5c). This was also the case for the CMG2 ligand present in serum (so far uncharacterized) ^20^, as well as for Collagen VI (Extended Data Fig. 4ab).

**Figure 5:**
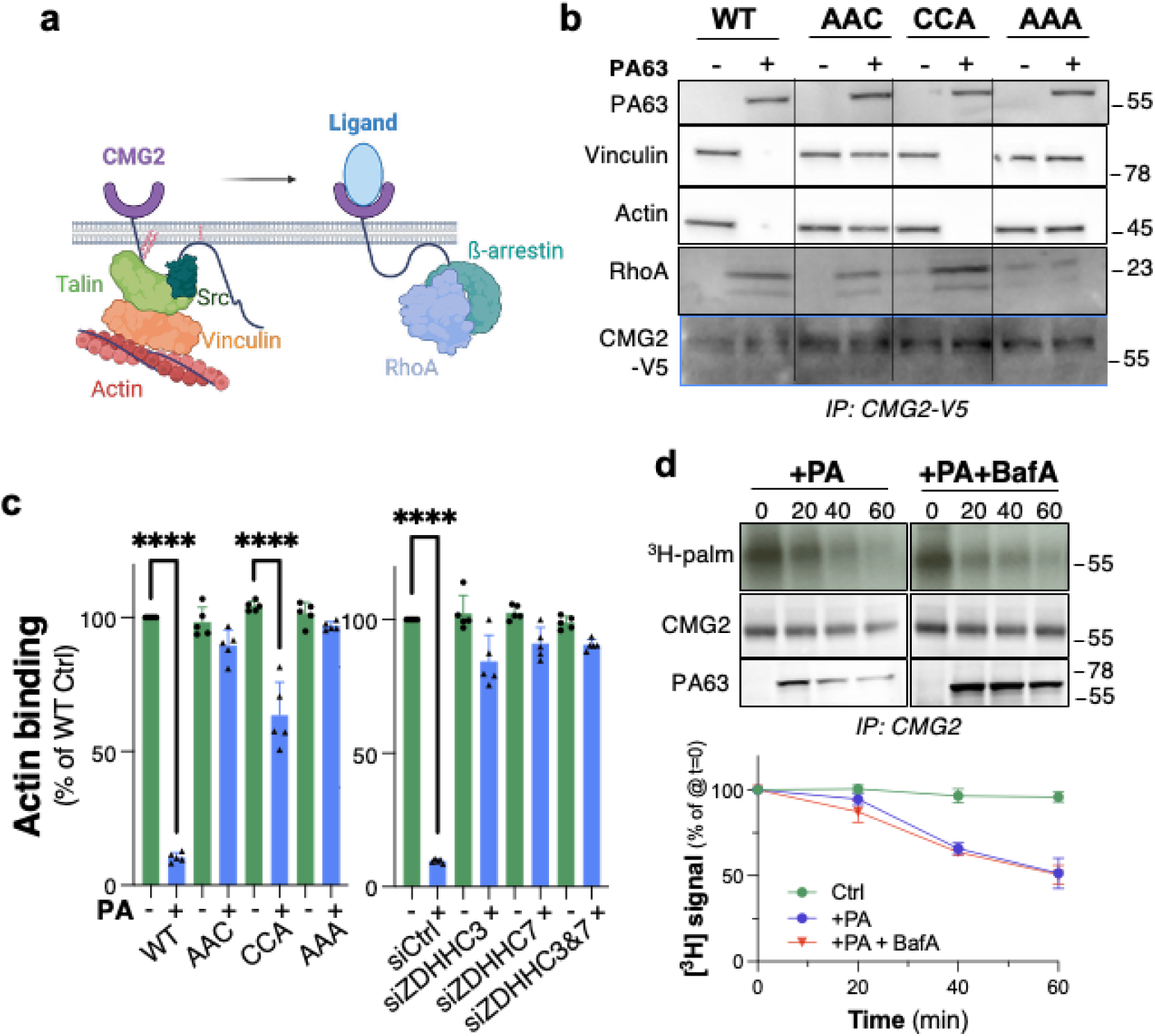
S-acylation controls the CMG2 TVA to RhoA signaling switch and ligand-induced deacylation. **a.** Schematic of CMG2 signalling at the plasma membrane. In the resting state CMG2 associates with Talin-Vinculin-Actin (TVA). Ligand binding (PA) disrupts TVA and promotes recruitment of RhoA and β-arrestin, enabling endocytosis and downstream cytoskeletal signalling. **b.** HeLa cells were transfected for 48 h with CMG2-V5 WT or cysteine mutants, then incubated ± PA (500 µg/ml; 1 h at 4 °C followed by 30 min at 37 °C). CMG2 was immunoprecipitated in PBS/1% Triton X-100 with anti-V5 for 4 h at 4 °C and analysed by SDS-PAGE and immunoblotting using anti-PA, anti-Talin, anti-Vinculin, anti-Actin, anti-RhoA, and anti-V5. **c.** Quantification of CMG2 association with Actin and APT2 from (b) for WT and cysteine mutants, and after 72 h siZDHHC3+siZDHHC7. **d.** RPE1 cells were labelled 2 h with [³H]-palmitate at 37 °C, washed into chase medium for 2.5 h, then treated ± PA (500 µg/ml) ± Bafilomycin A (100 nM). CMG2 was immunoprecipitated with anti-CMG2, analysed by SDS-PAGE and autoradiography (³H-palm) and immunoblotting with anti-CMG2 and anti-PA. Radiolabel incorporation was quantified by Typhoon; [³H]-palmitate at 2.5 h (pre-chase) was set to 100%, and timepoints were expressed relative to this (n=3, mean ± SD).

**Extended Data Figure 4:**
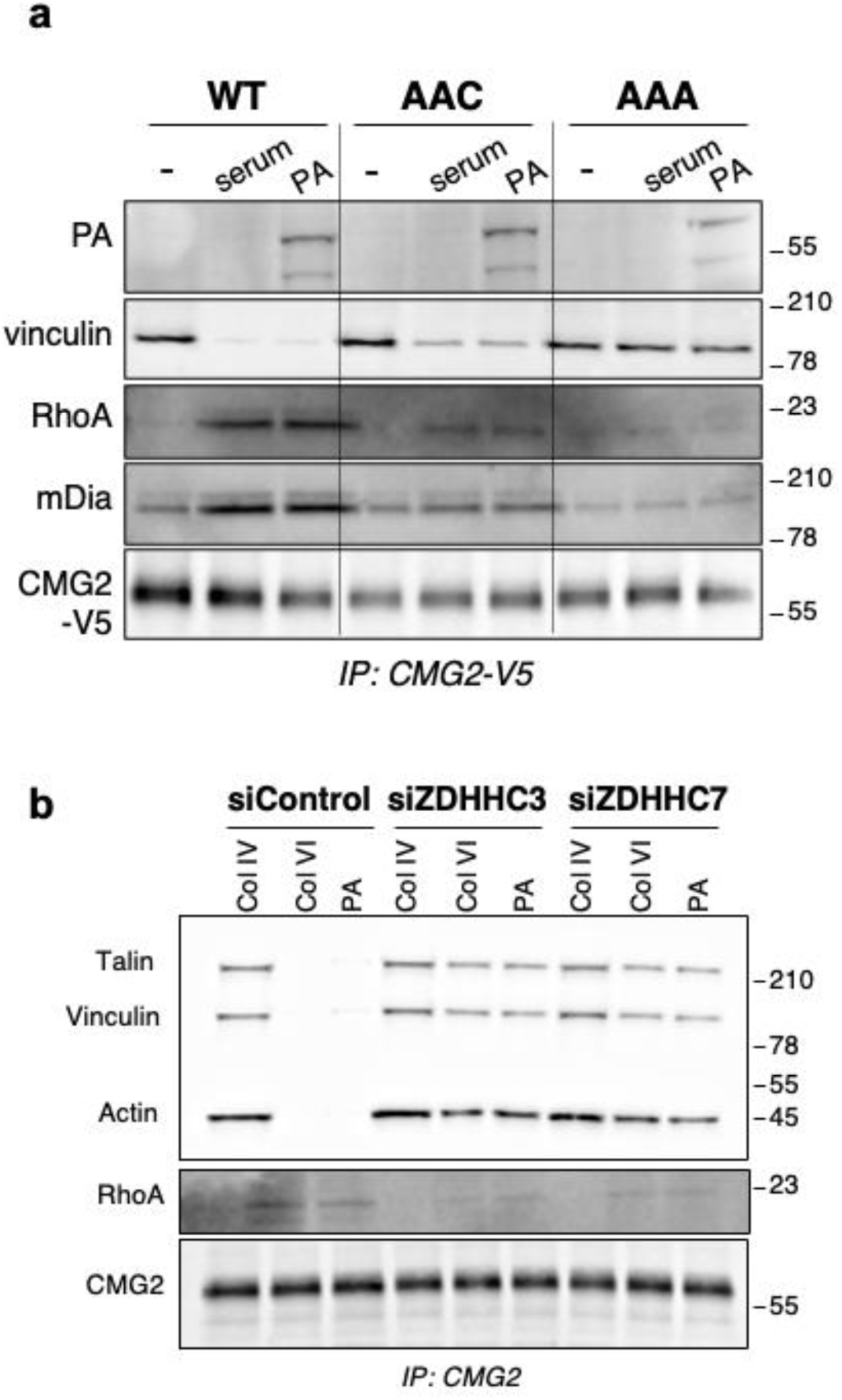
CMG2 ligands trigger the TVA to RhoA signalling switch. **a.** HeLa cells were transfected for 48 h with CMG2-V5 WT or cysteine mutants, switched to serum-free medium 24 h prior, and then incubated ± PA (500 µg/ml; 1 h at 4 °C followed by 30 min at 37 °C) or with 10% serum. CMG2-V5 was immunoprecipitated in PBS/1% Triton X-100 using anti-V5 and protein G for 4 h at 4 °C, resolved by SDS-PAGE, and blotted with anti-PA, anti-Talin, anti-Vinculin, anti-Actin, anti-RhoA, anti-mDia and anti-V5. **b.** RPE1 cells were silenced for 72 h with siZDHHC3, siZDHHC7 or control RNAi, switched to serum-free medium 24 h prior, and stimulated ± PA (500 µg/ml; 1 h at 4 °C followed by 30 min at 37 °C) or with collagen IV or VI (1 µg/ml; 3 h at 37 °C). CMG2 was immunoprecipitated in PBS/1% Triton X-100 using anti-CMG2 and protein G for 4 h at 4 °C, resolved by SDS–PAGE, and analysed by immunoblotting with anti-Talin, anti-Vinculin, anti-Actin, anti-RHOA and anti-CMG2-V5.

These findings show that CMG2 must be S-acylated for it to transduce ligand binding information into cytosolic signalling, likely by constraining the cytoplasmic tail so that the ligand-induced conformational change in the ectodomain can be propagated across the membrane, enabling release of the TVA complex.

### Ligand-induced CMG2 deacylation

We have previously shown that non-acylated anthrax toxin receptors lead to premature unproductive toxin uptake, i.e. where monomeric PA is internalized but the enzymatic subunits of the toxin remaining outside of the cell ^11^. This greater endocytic ability of non-acylated CMG2 led us to ask whether ligand binding could induce deacylation of WT CMG2. To test this, cells were labelled with [³H]-palmitate for 2 h, chased to allow CMG2 to reach the plasma membrane, and then stimulated with toxin. PA induced a time-dependent loss of [³H]-palmitate (Fig. 5d). This decay could not be prevented by treating cells with the vacuolar ATPase inhibitor Bafilomycin A, excluding the possibility that the observed loss of [³H]-palmitate occurred by lysosomal degradation of CMG2 (Fig. 5d). Thus, extracellular ligand binding triggers intracellular CMG2 deacylation, a regulatory step that we had previously hypothesized but could not directly demonstrate without the mechanistic insight into CMG2 acylation provided by the present study.

### Ligand binding triggers recruitment of membrane-bound APT2

We next tested whether ligand-induced CMG2 deacylation is mediated by APT2. We first tested whether the APT2 inhibitor ML349 would influence the TVA-to-RhoA switch. As revealed by coimmunoprecipitation of CMG2 with Talin, vinculin and actin but not with RhoA, ML349 fully blocked the switch for endogenous (Fig. 6a) and ectopically expressed CMG2 (Fig. 6b), the CCA and AAC mutants (Fig. 6b). The requirement for APT2 was further confirmed by APT2 silencing (Fig. 6c, Extended Data Fig. 5a) and knockout (Extended Data Fig. 5b). APT2 recruitment to CMG2 could be detected by immunoprecipitation of PA-treated cells (Fig. 6d). APT2 recruitment does not require CMG2 clustering following PA oligomerization since a PA mutant that cannot be processed by furin and therefore cannot oligomerize ^28,29^, still induced APT2 recruitment (Fig. 6d). APT2 recruitment was likewise independent of CMG2 interaction with the actin cytoskeleton, as the CMG2 W396L mutant, which cannot bind Talin ^20^, still recruited APT2 upon PA stimulation (Extended Data Fig. 5c).

**Figure 6:**
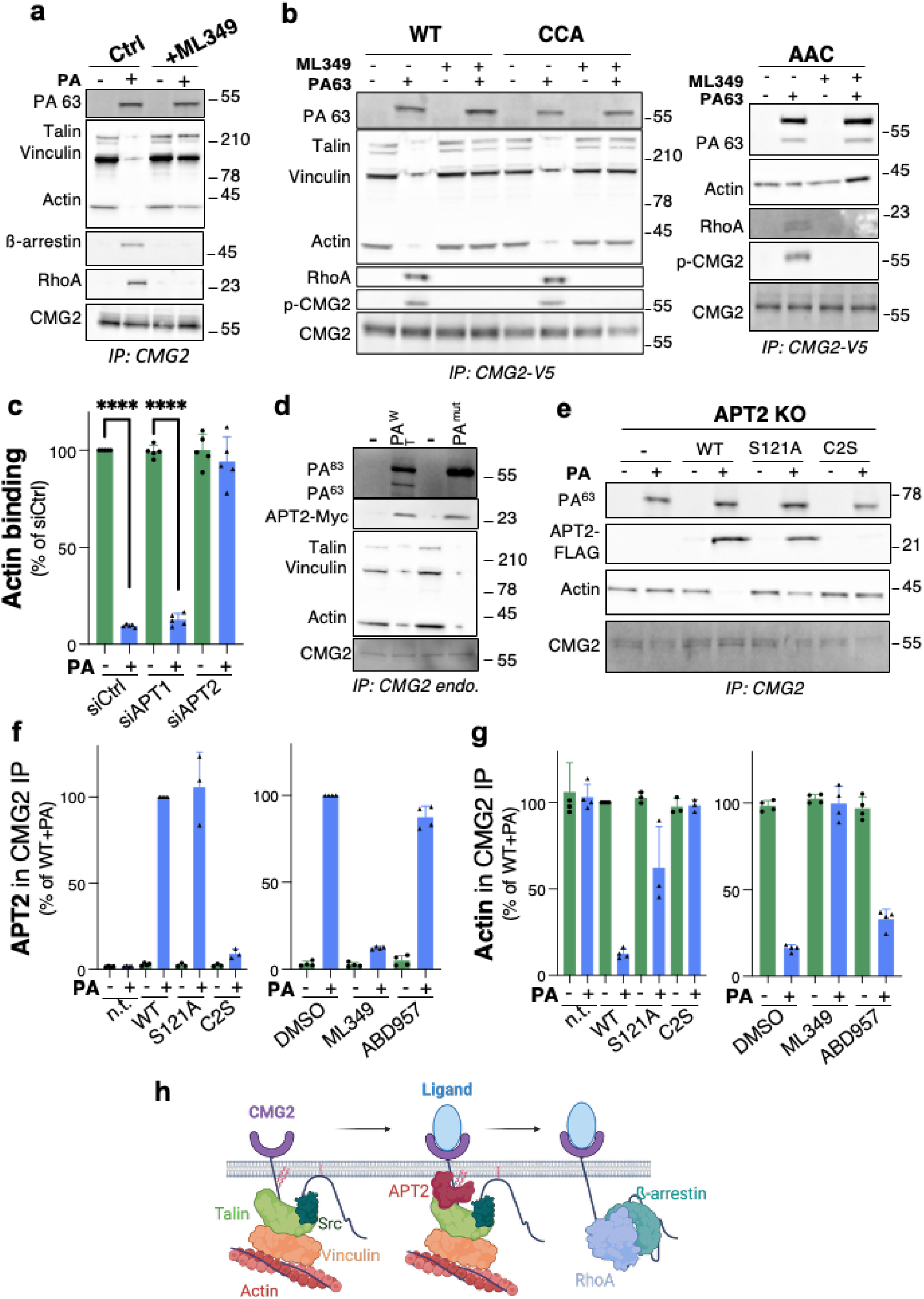
Ligand binding triggers APT2 recruitment and CMG2 deacylation. **a.** RPE1 cells were serum-starved, pre-incubated ± ML349 (APT2 inhibitor), and stimulated ± PA/LF at 4 °C followed by 37 °C; CMG2 was immunoprecipitated and analysed by SDS–PAGE and immunoblotting for PA, Talin, Vinculin, Actin, β-arrestin, RHOA and CMG2. **b.** HeLa cells were transfected 48 h with WT, CCA or AAC CMG2-V5, serum-starved, pre-treated ± ML349 and stimulated ± PA/LF; CMG2-V5 immunoprecipitates were analysed as in (a) and additionally probed for phosphotyrosine. **c.** RPE1 cells were silenced 3 d with siAPT1 or siAPT2, serum-starved and stimulated ± PA/LF; CMG2 immunoprecipitates were analysed for PA, Actin and CMG2, with Actin co-IP in control −PA normalized to 100% (n = 5, mean ± SD). **d.** RPE1 cells expressing APT2-Myc WT were stimulated with WT or uncleavable PA together with LF, and CMG2 immunoprecipitates were probed for PA, APT2-Myc, Talin, Vinculin, Actin and CMG2. **e,f,g.** APT2-KO RPE1 cells were rescued with APT2-FLAG WT, S121A, C2S or empty vector, serum-starved and stimulated with PA/LF; CMG2 immunoprecipitates were probed for PA, APT2-FLAG, Actin and CMG2. Quantification shows APT2 and Actin co-IP normalized to APT2-WT +PA = 100% (n = 3, mean ± SD). RPE1 cells pre-treated 4 h with DMSO, ML349 or ABD957 were analysed in parallel and expressed relative to DMSO +PA (n = 4, mean ± SD). **h.** Model of the CMG2 signalling switch in which ligand binding induces APT2 recruitment and CMG2 deacylation, releasing the Talin–Vinculin–Actin complex to permit endocytosis.

**Extended Data Figure 5:**
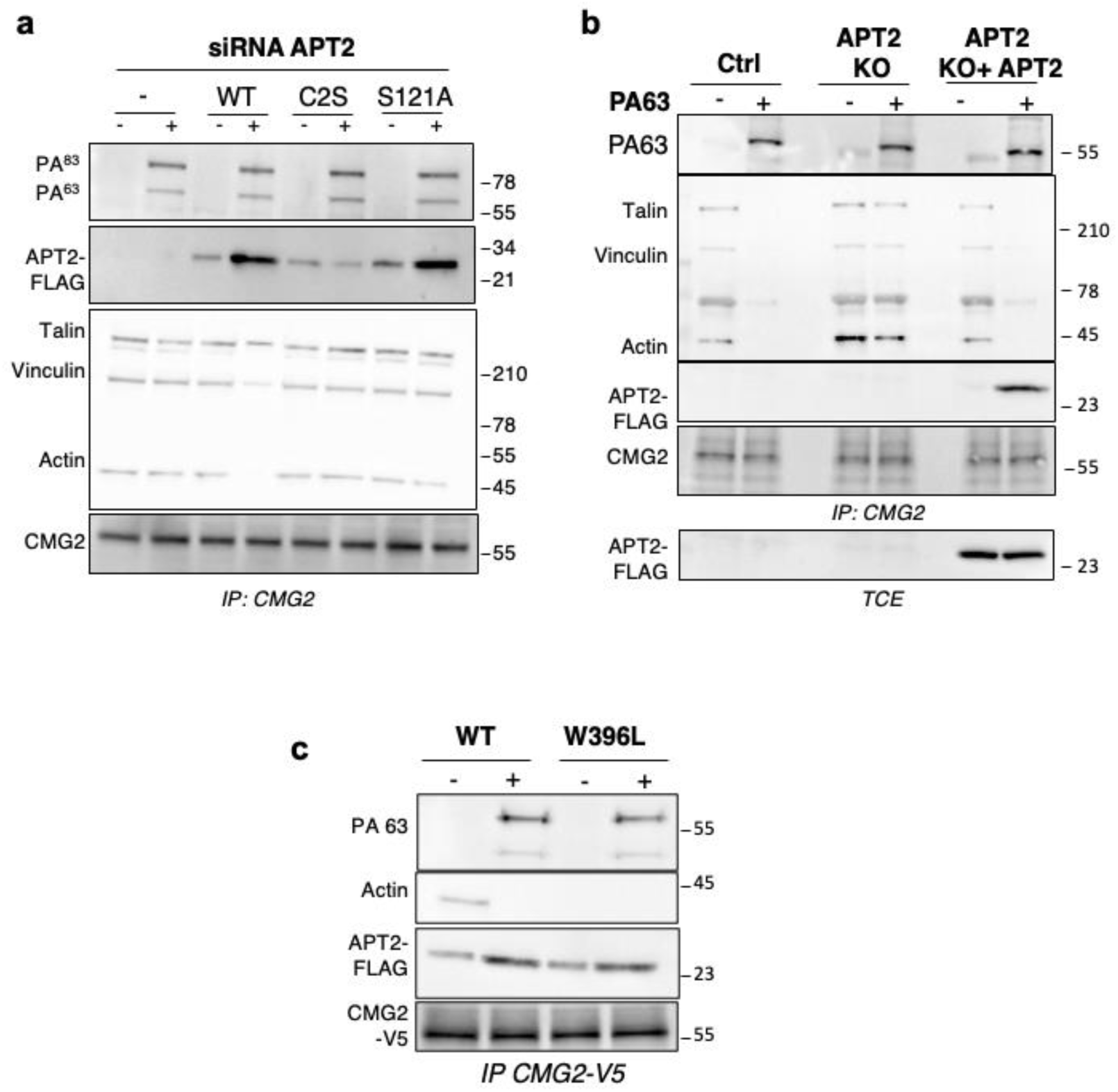
Ligand binding triggers recruitment of membrane-anchored APT2 to CMG2. **a.** RPE1 cells were silenced for 3 days with APT2 RNAi and transfected during the last 24 h with APT2-FLAG WT, C2S or S121A. Cells were serum-starved the day before, incubated or not for 1 h at 4 °C with PA (500 µg/ml) and LF (50 ng/ml), immunoprecipitated in PBS–1% TX-100 with anti-CMG2 plus protein G for 4 h at 4 °C, and analysed by SDS–PAGE and immunoblotting using anti-PA, anti-APT2-FLAG, anti-Talin, anti-Vinculin, anti-Actin and anti-CMG2 antibodies. **b.** WT or APT2-KO RPE1 cells were transfected for 24 h with APT2-FLAG WT, serum-starved, incubated or not with PA/LF under the same conditions, and CMG2 immunoprecipitates were analysed by SDS–PAGE and immunoblotting for PA, APT2-FLAG, Talin, Vinculin, Actin and CMG2; total cell extracts were probed to verify APT2-FLAG expression. **c.** HeLa cells were transfected for 48 h with WT or W396L CMG2-V5 together with APT2-FLAG, serum-starved overnight, incubated or not for 1 h at 4 °C plus 30 min at 37 °C with PA (500 µg/ml) and LF (50 ng/ml), immunoprecipitated with anti-CMG2-V5 overnight at 4 °C, and analysed by SDS-PAGE and immunoblotting with anti-PA, anti-APT2-FLAG, anti-Actin and anti-CMG2-V5.

APT2 is a soluble globular protein that can interact with membranes following a three-step process involving electrostatic interactions, insertion of a hydrophobic loop and S-acylation on Cys2 ^22^. The acylation-deficient C2S mutant was unable to bind CMG2 (Fig. 6ef), confirming that APT2 must itself be S-acylated to deacylate its substrates ^22^. The catalytically inactive S121A APT2 mutant in contrast was able to bind CMG2, but failed to trigger TVA release (Fig. 6ef), confirming the requirement for the thioesterase activity. Interestingly, ML349 treatment prevented APT2 recruitment (Fig. 6f). Since ML349 inserts into the hydrophobic acyl-chain binding pocket of APT2 ^30^, these results indicate that for APT2-CMG2 interactions to be detected by immunoprecipitation, a CMG2-bound acyl chain must insert into the hydrophobic pocket of membrane-anchored APT2. APT2 recruitment always coincided with loss of binding of the actin cytoskeleton to CMG2 (Fig. 6g).

Thus, recruitment of membrane-anchored APT2 constitutes the first cytosolic response to ligand binding, and deacylation acts as the execution step of the CMG2 signalling switch, converting a cytoskeleton-anchored receptor into an internalization-competent complex (Fig. 6h).

### APT2 inhibition prevents anthrax intoxication

We next examined the role of APT2-mediated deacylation in anthrax toxin entry. The mode of action of anthrax lethal toxin, namely the combination of Protective Antigen PA with lethal Factor (LF), involves binding of the PA63 form to CMG2/TEM8, formation of the PA oligomer to which LF can bind, endocytosis, conversion of the PA oligomer into a PA pore, which is required for LF to cross endosomal membranes and ultimately reach the cytosol ^31,32^. In the cytosol, LF eventually cleaves MAP kinase kinases such as MEK1 or MEK2. We first tested the role of APT2 in anthrax toxin intoxication by monitoring MEK2 levels by immunofluorescence ^12^. In this assay, control cells stain positive with an antibody directed to the N-terminus of MEK2, while lethal toxin treated cells are negative, LF cleaving MEK2 ^33^. In this assay MEK2 remained readily detectable in APT2 KO cells treated with lethal toxin (Fig. 7ab), the differences in MEK2 levels in the untreated cells hindered the analysis.

We therefore monitored the different steps of lethal toxin’s mode of action biochemically. Treatment with the APT2 inhibitor ML349, but not the APT1 inhibitor ML348, drastically inhibited PA pore formation (Fig. 7cd). This effect was not due to impaired PA oligomerization, as acidification of cell lysates prior to SDS–PAGE ^21^ revealed normal oligomer assembly (Fig. 7e, Extended Data 6a). Consistently with impaired PA pore formation, MEK2 cleavage was not detected in ML349-treated cells (Fig. 7c). In further agreement, APT2 silencing (Extended Data 6b) and APT2 knockout (Extended Data 6c) strongly reduced PA pore formation and MAPK kinase cleavage, and both phenotypes were rescued by re-expression of APT2 (Extended Data Fig. 6d). Importantly, APT2 knockout did not cause a general defect in endocytosis or in the endo-lysosomal trafficking pathway, as assessed by the normal EGF internalization and degradation observed in APT2 KO cells (Extended Data Fig. 7). APT2-mediated deacylation was also required for the physiological function of CMG2, as ML349 treatment impaired collagen VI degradation (Fig. 7g).

**Figure 7:**
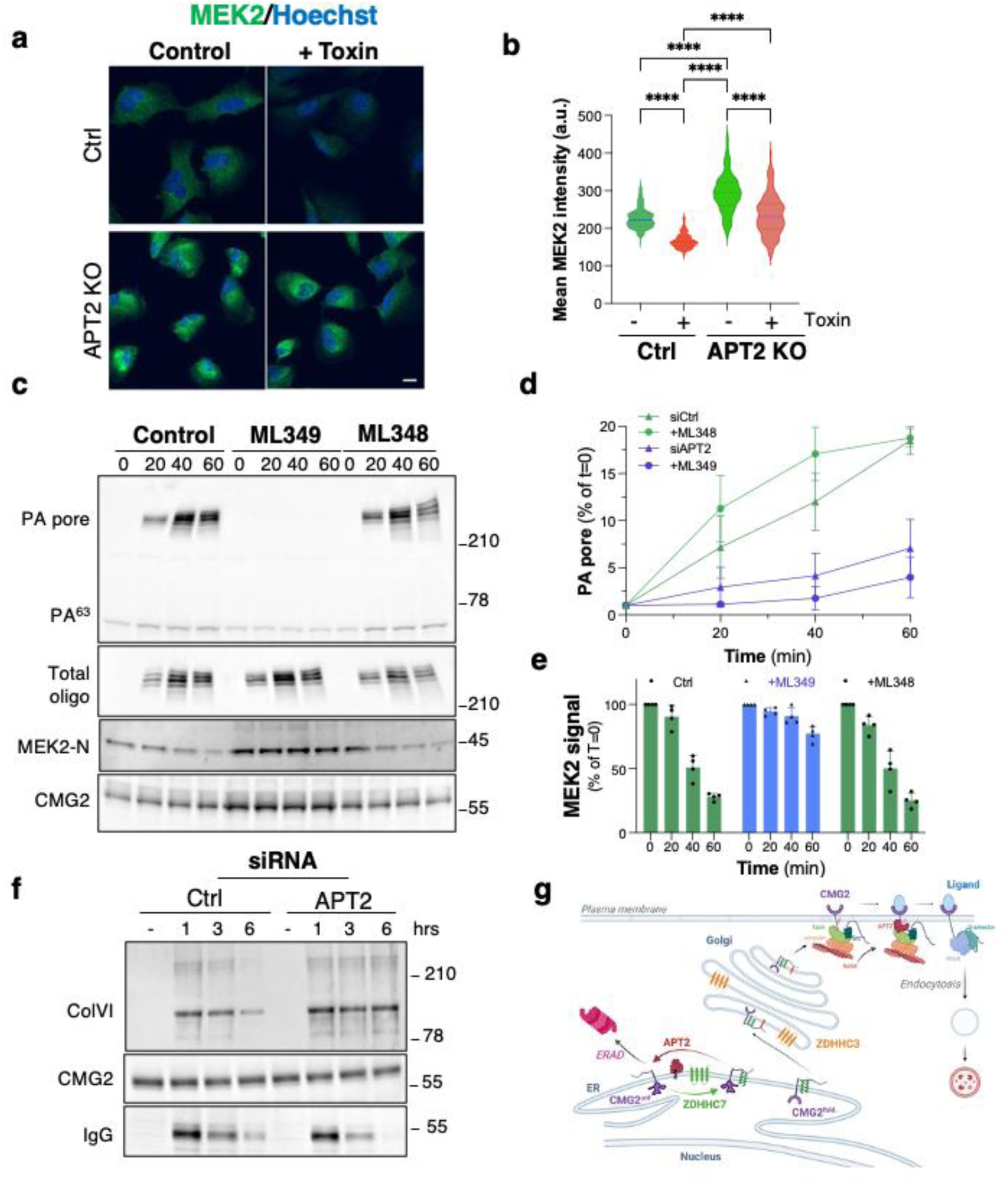
CMG2 deacylation is essential for endocytosis of Anthrax lethal toxin. **a.** Representative confocal images ofRPE1 cells, WT or knocked for APT2. Cells were incubated 1 hour at 4°C with 500 ng/ml Anthrax toxin Protective Antigen (PA) and 100ng/ml Lethal Factor LF; cells were washed and incubated for 2h30 at 37°C. Mek2 was stained by immunofluorescence, and nuclei were stained with Hoechst. Scale bar = 10μm. **b.** Quantification of **a**. Cells were segmented using Cellpose3 and Mek2 mean staining intensity was measured for each cell. Groupped were compared using a Kruskal-Wallis test, ***: p-value < 0.0001. **c**. RPE1 cells were pre-incubated 4 hours at 37°C with 10µg/ml ML348 or ML349. Cells were incubated 1 hour at 4°C with 500µg/ml PA and 50ng/ml LF and chased different indicated times at 37°C. 40 µg of protein samples were submitted to SDS-PAGE, and analyzed by immunoblotting with rabbit anti-human CMG2, rabbit anti-PA or rabbit anti-MEK2 N-terminal. The amount of PA pore (SDS-resistant) (**d**) and cleaved MEK2N-terminal (**e**) were quantified, and set to 1 after 1 hour at 4°C before chase, and different time of chase were expressed relative to this (n=4, error bars represent standard deviation). **f.** RPE1 cells were silenced for 3 days with human APT2 RNAi or control RNAi. Cells were changed 24h prior experiment in serum free medium, incubated with 1ug/ml Collagen6 indicated hours. 40 °g of protein samples were submitted to SDS-PAGE, and analyzed by immunoblotting with rabbit anti-collagen VI, anti-CMG2, and anti-IgG rabbit. **g.** Cartoon of the S-acylation and deacylation events during the life cycle of CMG2.

**Extended Figure 6:**
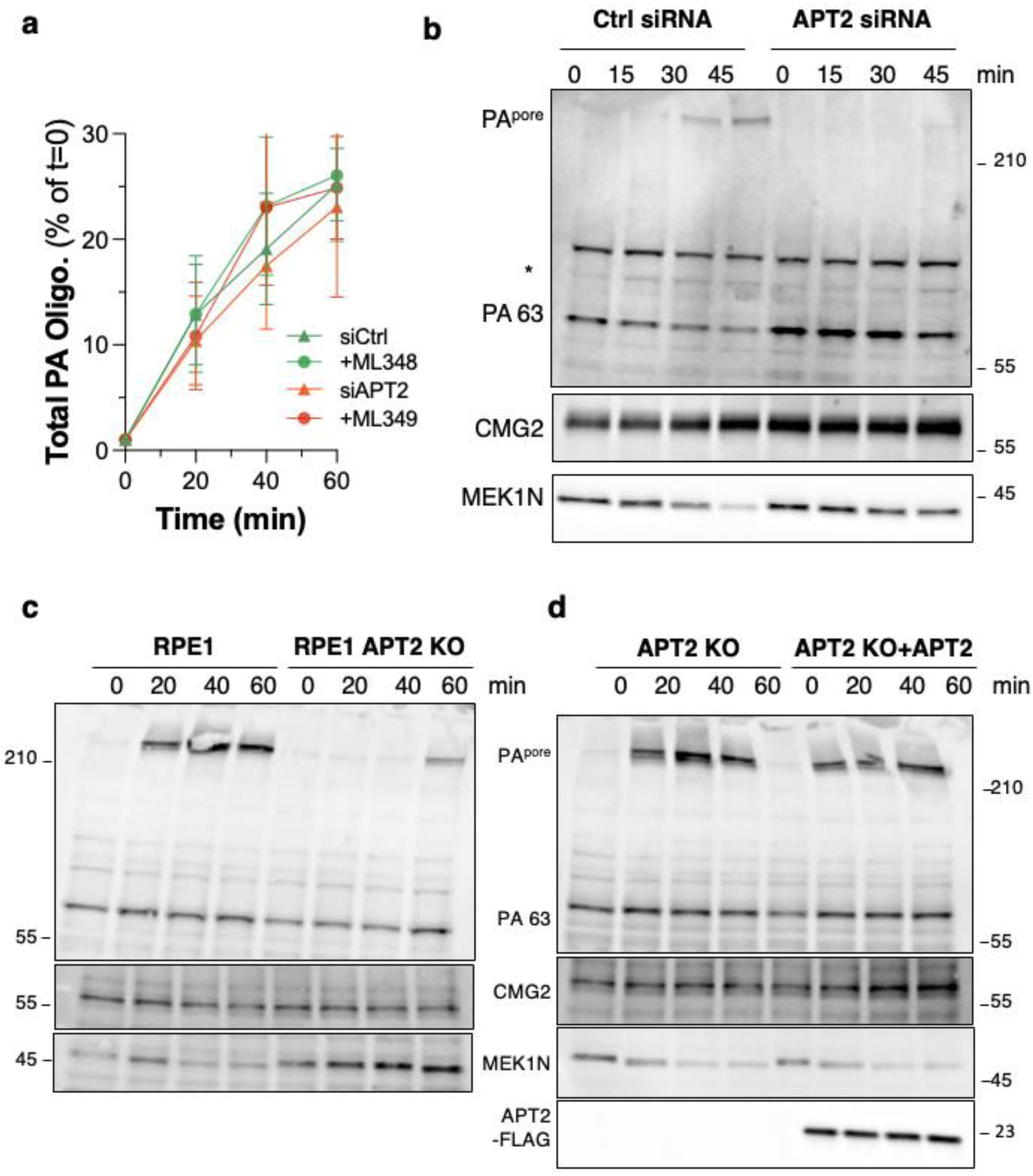
RPE1 cells were pre-incubated 4 hours at 37°C with 10µg/ml ML348 or ML349 or were silence for APT2 expression during 72 hrs. Cells were incubated 1 hour at 4°C with 500µg/ml PA and 50ng/ml LF and chased different indicated times at 37°C. **a**. 40 µg of protein samples were acidified to convert all PA oligomers to the SDS resistant form, submitted to SDS-PAGE, and analyzed by immunoblotting with rabbit anti-PA or rabbit anti-MEK2 N-terminal. The amounts of PA oligomers were quantified, and plotted (n=4, error bars represent standard deviation). **b**. 40 µg of protein samples were submitted to SDS-PAGE, and analyzed by immunoblotting with rabbit anti-human CMG2, rabbit anti-PA or rabbit anti-MEK2 N-terminal. **c**, **d**. Control and APT2 RPE1 were treated and analyzed as in ab.

**Extended Figure 7:**
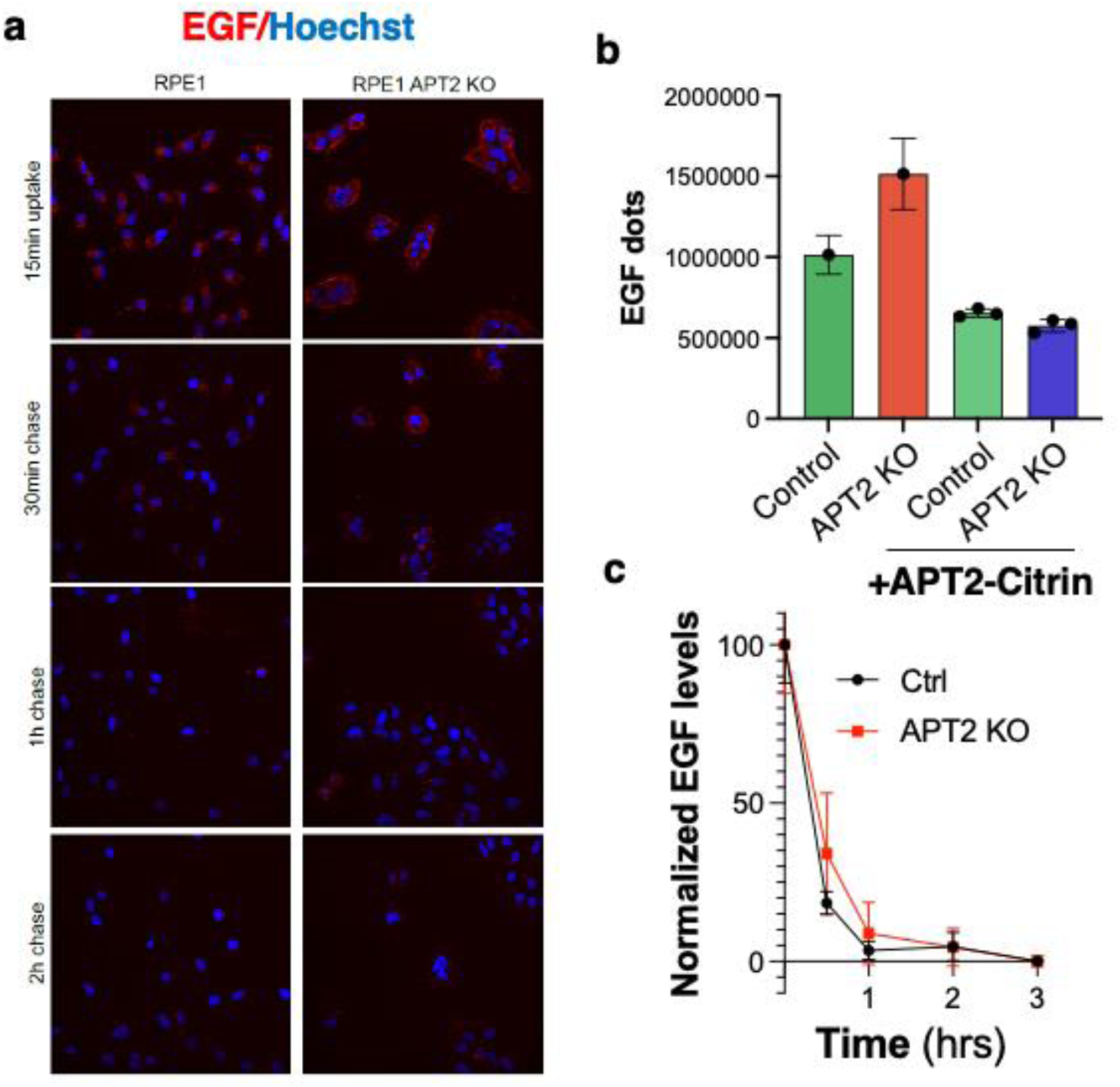
EGF uptake and endo-lysosomal trafficking are not impaired in APT2 KO cells. **a**. Spinning disk automated microscope confocal images showing RPE1 and RPE1 APT2 KO cells incubated during 15min at 37°C with 0.5 μg/ml A647-EGF and chased for the specified times. b. Graph showing the integrated intensity sum of EGF dots per cell after 15 min of A647-EGF uptake in RPE1 vs RPE1 APT2 KO transfected or not with APT2 citrin. c. Graph showing the normalized EGF dots integrated intensity sum per cell at the different timepoints of the pulse chase experiment in both Control and APT2 KO cells transfected. For each data point on the different graphs, a minimum of 1000 cells were quantified. Data represents the mean ±SD.

Together, these findings demonstrate that APT2-mediated deacylation is essential for CMG2 to transition into an internalization-competent state, and is therefore essential for anthrax toxin entry into cells and collagen VI homeostasis.

## DISCUSSION

CMG2 is a multifunctional receptor that links extracellular matrix homeostasis to cytoskeletal signalling and endocytosis, and is also hijacked as the primary host receptor by anthrax toxin. Here we show that CMG2 biogenesis and function are controlled by sequential S-acylation–deacylation cycles acting across the secretory and endocytic pathways. These cycles enable CMG2 to adopt distinct conformational or interaction states that correspond to discrete stages of its lifecycle: folding in the ER, transport to the plasma membrane, and ligand-induced signalling and internalization. The lipidation events we describe therefore couple CMG2 biogenesis with its function as a signalling and endocytic receptor at the plasma membrane.

Our findings demonstrate that S-acylation of the juxtamembrane cysteines by ZDHHC7 in the ER promotes productive folding by preventing premature degradation of newly synthesized CMG2 by ERAD. Conversely, APT2-mediated removal of these acyl chains renders folding intermediates vulnerable to ERAD. Thus, acute APT2 inhibition increases CMG2 levels in cells and in vivo. These observations establish S-acylation as a key positive determinant of CMG2 protein homeostasis. Since a subset of Hyaline Fibromatosis Syndrome-associated missense mutations result in impaired CMG2 folding kinetics, chronic partial APT2 inhibition represents a rational therapeutic strategy to increase CMG2 expression in these patients. This mechanism is conceptually related to pharmacological strategies aimed at stabilizing CFTR and other folding-compromised mutated transmembrane proteins.

After ER exit, a second acylation event, mediated by ZDHHC3 in the Golgi on Cys379, allows CMG2 to interact with Arf6 and traffic to the plasma membrane. This indicates that CMG2 biogenesis is regulated not only at the folding stage but also during transport through the secretory pathway, with distinct S-acylation events conferring sequential benefits.

At the cell surface, we find that CMG2 must be S-acylated for the ligand-induced release of from TVA and the recruitment of RhoA and β-arrestin to occur. These data support a model in which S-acylation constrains the cytoplasmic tail in a conformation that is competent to propagate the ligand-induced conformational change across the transmembrane helix, allowing CMG2 to reposition its cytosolic interactions. In the absence of S-acylation, ligand binding still occurs but the conformational signal fails to cross the membrane, preventing receptor activation.

Finally, we show that ligand binding triggers the recruitment of APT2 to CMG2, involving the insertion of a CMG2-bound acyl chain into the hydrophobic acyl-chain binding pocket of APT2. APT2 recruitment precedes TVA release, and mediates CMG2 deacylation, which converts the receptor from a cytoskeleton-anchored to actin regulatory and internalization-competent state. Genetic or pharmacological inhibition of APT2 therefore blocks anthrax lethal toxin internalization, pore formation in endosomes, and downstream MAPK cleavage in the cytosol. Similarly, in the physiological setting, APT2 inhibition prevents CMG2-mediated collagen VI degradation. These results demonstrate that deacylation is not simply the reversal of a modification that favours biogenesis, but a signalling execution step. Treatment of an anthrax infection would require an acute and maximal inhibition of APT2 activity, allowing the immune system to clear the infection. In contrast, HFS treatment would require a chronic low concentration or intermittent treatment, allowing an increase in CMG2 protein expression, but at a dose that would only mildly or transiently affect CMG2 signalling ability and ECM regulation.

In summary, this work identifies coordinated and temporally ordered S-acylation–deacylation cycles as a central regulatory principle controlling CMG2 folding, trafficking, and function, both physiological and pathological. Hence, this study demonstrates that S-acylation–deacylation cycles can control sequential trafficking steps along the biosynthetic and endocytic pathway. Beyond clarifying the molecular logic of CMG2 acylation, our findings establish APT2 as a druggable regulator of receptor function, with implications for both HFS therapy and anthrax infection, as well as other genetic diseases affecting the folding of membrane proteins, 50to 70% of which undergo S-acylation^34–37^.

## Acknowledgments

We warmly thank Arthur Samurkas for generating Fig. 1a, all the members of the F.G.v.d.G. lab for their discussions, suggestions and critical reading of the manuscript, the staff of the ACCESS-Geneva biomolecular screening platform for the high content image analysis, the staff of the Center for Phenogenomics of the EPFL and the staff of the EPFL Bio Imaging core facility. We thank Prof. Paolo Bonaldo for providing us with purified Collagen VI tetramer, and Prof Jeff Brodsky for enlightening discussions that have contributed to the conceptual advances provided by this work. This work was supported by the Swiss National Science Foundation (310030-214874 and 3200-0-239873), by the Gelù Foundation, and by generous donors represented by CARIGEST SA, such as Fondazione Teofilo Rossi di Montelera e di Premuda.

## Author contributions

Conceptualization, L.A., S. B., O. J., F.S.M., G.V.D.G.; Investigation, L.A., S. B., O. J., F. S. M., B. K., V. M.; Funding Acquisition, G.V.D.G.; Writing–Original Draft L.A., G.V.D.G.; Writing–Review & Editing, L.A., S. B., O. J., F. S. M., V. M., G.V.D.G.; Resources, L.A., S.B., B. K.

## Declaration of Interests

The authors declare no competing interests.

## MATERIALS AND METHODS

### Cell lines

HeLa cells (ATCC CCL-2) were grown in Modified Eagle’s Medium (Sigma Life Science) supplemented with 10% FBS, 2 mM of L-glutamine, non-essential amino acids, penicillin, and streptomycin (GIBCO). Primary fibroblasts and RPE1 (ATCC CRL-4000) and RPE1 KO APT2 ^22^ were grown in Dulbecco’s Modified Eagle Medium (Sigma Life Science) supplemented with 10% FBS, penicillin, and streptomycin (GIBCO).

### Plasmids and transfection

Human CMG2 (isoform 4, UniProt P58335-4) WT was cloned in a pcDNA3.5/V5-HIS-TOPO expression vector or in pHS003-EGFP. Cystein mutants and W396L mutant were generated in pcDNA3.5/V5-HIS-TOPO using the QuikChange Site-directed Mutagenesis Kit (Agilent) according to the manufacturer’s protocol.

Human APT2/LYPLA2 WT was cloned in a pCMV6-myc-flag expression vector. S121A and C2S APT2 mutants were generated in pCMV6-myc-flag using QuikChange Site-directed Mutagenesis Kit (Agilent) according to the manufacturer’s protocol.

MYC fusion human ZDHHC3 and human ZDHHC7 were cloned in pcDNA3.1^38^. Plasmids were transfected into cells for 24 h or 48 h, depending on the experiment, using *TRANS*IT–X2 according to manufacturer’s protocol (MIRUS).

Verified siRNA for human ZDHHCs and for human APT1 and APT2 were purchased from Qiagen. APT1 target sequence: 5’-AACAAACTTATGGGTAATAAA-3’. APT2 target sequence: 5’-CAGCTGCTTCTCAGTCATGAA-3’. ARF6 target sequence: 5’-TGCGACCACTATGATAATATT-3’. The pool of siRNA were done as followed: Mix1 (ZDHHC1 (5’-ACCGGCTGTGATGCTCCAATA-3’) + ZDHHC3 (5’-TCCGTTCTCATGAATGTTTAA-3’) + ZDHHC7 (5’-CCCGTGGTTACTATGAATGTA-3’) + ZDHHC13 (5’-CAGCATAGTAGCCTTTCTATA-3’) + ZDHHC17 (5’-CAGTACCTGTTTGATACGAAA-3’); Mix2 (ZDHHC2 (5’-TAGCTACTGCTAGAAGTCTTA-3’) + ZDHHC6 (5’-GAGGTTTACGATACTGGTTAT-3’) + ZDHHC4 (5’-ATGGATTGCTTCATTACCTTT-3’) + ZDHHC16 (5’-CTCGGGTGCTCTTACCTTCTA-3’)); Mix3 (ZDHHC5 (5’-ACCACCATTGCCAGACTACAA-3’)+ZDHHC8(5’- CCGGGCTCCGCTGTACAAGAA-3’) + ZDHHC15 (5’-ATCGCTATATCAAGTATCTAA-3’) + ZDHHC20 (5’- TACCTGTTATGAGTTGCCTATA-3’)); Mix4 (ZDHHC9 (5’- CTCAACCAGACAACCAATGAA-3’) + ZDHHC12 (5’- CAGATACTGCCTGGTGCTGCA-3’) + ZDHHC18 (5’- AAGCCTGATGCCAGCATGGTA-3’)); Mix5 (ZDHHC11 (5’- CGCGTGGAAATACATTGCCTA-3’) + ZDHHC23 (5’- CTGCGAGTACATAGATCGGAA-3’) + ZDHHC24 (5’- CCGCTGCGTGGGGCTTCGGCAA-3’); Mix6 (ZDHHC14 (5’- CTGGGTGTCCTCGGCAAAGTT-3’) +ZDHHC19 (5’- GACCCTGGCATCTTACATCAA-3’) + ZDHHC21 (5’- GTGGGACTAATACAAGATCTA-3’) +ZDHHC22 (5’-TGGGTTCATTTATGCCCTATA-3’).

As a control siRNA, a sequence targeting the viral glycoprotein VSV-G (5’-ATTGAACAAACGAAACAAGGA-3’) was used. Transfections of 50 nM of siRNA were carried out using *TRANS*IT-X2 (MIRUS), and the cells were analyzed at least 72 h after transfection.

### Antibodies and Reagents

Anti-mouse CMG2 monoclonal antibody 8F7 was generated by the genetic immunization of rats with a mouse CMG2 construct^39^ (Genovac, Germany). The other antibodies were commercially available: goat anti-human CMG2 (R&D, AF2940, RRID: AB_2056740); rabbit anti-human CMG2 (Proteintech, RRID: AB_2056741); goat anti-human TEM8 (R&D, ref: AF3886, RRID: AB_2056726); rabbit anti-ARF6 (LSBio, ref: LS-C81881,RRID: AB_2058487); mouse anti-TRAP alpha (Santa Cruz, RRID: AB_1130631); anti-V5 (Invitrogen, 46-0705, RRID: AB_2556564); anti-GFP (Takara Bio, RIDD: AB_11153295); mouse anti-Myc (SIGMA, 9E10 M4439, RRID: AB_439694), anti-human ZDHHC3 (Abcam, ab31837, RRID: AB_742236); anti-GM130 (BD biosciences, 610823, RRID:AB_398142); mouse anti-GAPDH (Acris Antibodies, 4A1-MA0100, RRID_AB1874646); anti-actin (Merck Millipore, RRID: AB_2223041); anti-Talin (Sigma, RRID: AB_477572); anti-Vinculin (Sigma, RRID: AB_477629); anti-phospho tyrosine (Millipore, RRID: AB_916371); anti-RhoA (Cell signalling, 67B9, RIDD: AB_10693922); anti-SRC (Abcam, ref: 16885, RRID: AB_443522); anti-Anthrax Protective Antigen (List Biological Laboratories, ref: 771B); anti-MEK2 (Santa Cruz, ref:12578, RRID: AB_648958), anti-collagen VI (Fitzgerald laboratories, ref: 70R-CR009X, RRID: AB_1283876). Anti-IgG (Santa Cruz, ref: SC-2025, RRID: AB_737182), HRP-conjugated secondary antibodies (Pierce); and for immunoprecipitation protein G beads were purchased from GE Healthcare.

Anti MEK1-N terminal antibodies were produced in our laboratory (Abrami et al., 2008).

Brefeldin A (Sigma) was used at 2ug/ml, Bafilomycin A (Sigma) was used at 100 nM and Cycloheximide (Sigma) was used 10ug/ml 1 hour before and during 3H-palmitic acid experiments. ABD957, ML348 and ML349 (Cayman Chemical company) were used at 10uM indicated times. Palmostatin B (Merk) was used at 50uM indicated times.

Wild type PA (Anthrax toxin Protective Antigen) and LF (Anthrax toxin Lethal Factor) were produced in our laboratory by overexpression in *E.coli* as described (Feld et al., 2012)^40^. Collagen VI was a gift from Bonaldo Laboratory^41^.

### Western Blotting

Cells were washed three times in 1x PBS at 4°C and lysed in Buffer (1x PBS, 1% Triton X-100, and protease inhibitor cocktail; Roche) for 30 min on ice. Lysates were then spun down at 5,000 rpm on a table-top centrifuge, and the protein content of supernatant were determined, the samples were boiled in Laemmli buffer for 5 min before separation via SDS-PAGE and western blotting against the different anti-bodies used in this study. Western blots were developed using the ECL protocol (Thermo Fisher) and imaged on a Fusion Solo from Vilber Lourmat. Densitometric analysis was performed using the software Bio-1D from the manufacturer.

### Acyl-RAC capture assay

Protein S-palmitoylation was assessed by the Acyl-RAC assay^42^ as previously described ^43^ with some modifications. Mice tissues were lysed in 400 μl buffer (0.5% Triton-X100, 25 mM HEPES, 25 mM NaCl, 1 mM EDTA, pH 7.4, and protease inhibitor cocktail). Then, 200 μl of blocking buffer (100 mM HEPES, 1 mM EDTA, 87.5 mM SDS, and 1.5% [v/v] methyl methanethiosulfonate [MMTS]) was added to the lysates and incubated for 4 h at 40°C to block free the SH groups with MMTS. Proteins were acetone precipitated and resuspended in buffer (100 mM HEPES, 1 mM EDTA, 35 mM SDS). For treatment with hydroxylamine (NH2OH) and capture by Thiopropyl Sepharose® beads, 2 M of hydroxylamine was added together with the beads (previously activated for 15 min with water) to a final concentration of 0.5 M of hydroxylamine and 10% (w/v) beads. As a negative control, 2 M Tris was used instead of hydroxylamine. These samples were then incubated overnight at room temperature on a rotating wheel. After washes, the proteins were eluted from the beads by incubation in 40 μl SDS sample buffer with ß-mercaptoethanol for 5 min at 95°C. Finally, samples were separated by SDS-PAGE and analyzed by immunoblotting. A fraction of the tissue lysate was saved as the input.

### ^3^H-palmitic acid radiolabeling experiments and immunoprecipitation

HeLa, RPE1, human fibroblast cells were transfected or not with different constructs, incubated for 2 h in Glasgow minimal essential medium (IM; buffered with 10 mM HEPES, pH 7.4) with 200 µCi/ml of ^3^H palmitic acid (9,10-^3^H[N]) (American Radiolabeled Chemicals, Inc.)^43^. The cells were washed, and incubated in DMEM complete medium for the indicated time of chase or directly lysed for immunoprecipitation with the indicated antibodies.

For immunoprecipitation, cells were washed three times in PBS, lysed 30 min at 4°C in the IP buffer (0.5% Nonidet P-40, 500 mM Tris pH 7.4, 20 mM EDTA, 10 mM NaF, 2 mM benzamidin and protease inhibitor cocktail [Roche]), and centrifuged 3 min at 5000 rpm. Supernatants were subjected to preclearing with Protein G Sepharose beads prior to the immunoprecipitation reaction. Supernatants were incubated overnight with the appropriate antibodies (V5 or goat anti-human CMG2) and Protein G Sepharose beads. After immunoprecipitation, washed beads were incubated for 5 min at 90°C in reducing sample buffer prior to 4−20% gradient SDS-PAGE. Gels were incubated 30 min in a fixative solution (25% isopropanol, 65% H2O, 10% acetic acid), followed by a 30 min incubation with the signal enhancer Amplify NAMP100 (GE Healthcare). The radiolabeled products were imaged using a Typhoon phosphoimager and quantified using a Typhoon Imager (ImageQuanTool, GE Healthcare). The images shown for ^3^H-palmitate labeling were obtained using fluorography on film.

### Immunofluorescence microscopy

Cells were fixed in 4% paraformaldehyde (15 min), quenched with 50 mM NH_4_Cl (30 min) and permeabilized with 0.1% Triton X-100 (5 min). Antibodies were diluted in PBS containing 1% BSA. Coverslips were incubated for 1 h with primary antibodies, washed three times in PBS, and incubated 45 min with secondary antibodies. Coverslips were mounted onto microscope slides with ProLong^TM^ Gold Antifade Mountant (P36930; Invitrogen). Images were collected using a confocal laser-scanning microscope (Zeiss LSM 700) and processed using Fiji™ software.

For automated microscopy, HeLa Cells were seeded at 5× 10^3^ cells per well in Ibidi 96-well plates, silencing was performed directly in presence of siRNAs in transfection mix containing *TRANS*IT-X2 (MIRUS). After 72 hours, the cells were fixed using 3% PFA, washed and subjected to immunofluorescence.

### EGF uptake and degradation by automated spinning disk confocal microscopy

RPE1 and RPE1 APT2 KO Cells were seeded at 1× 10^4^ cells per well in Ibidi 96-well plates. After 24 hours, cells were transfected using *TRANS*IT-X2 (MIRUS) with 0.1μg of APT2-citrin plasmid per well. 24 hours later, the medium was replaced by warm medium containing 0.5 μg/ml of A647-EGF and the cells were incubated 15min at 37°C, washed twice and further incubated in warm medium at 37°C for the specified times of the pulse chase experiment and finally fixed 15 minutes with 3% PFA^44^. After the fixation, the cells were incubated with PBS containing 0.01% saponin, 0.5% BSA and Hoechst diluted 2000 times. The plate was washed 3 times using a Biotek EL406 plate washer and the images acquired using a 40X water immersion objective with 1.15NA on a spinning disk confocal automated microscope (Molecular Device). For each FOV, 5 stacks separated by 1 μm were acquired and combined into a maximum intensity projection image. Analysis of the images was done via the MetaXpress Custom Module editor software. The images were segmented to generate relevant masks, which were then applied on the fluorescence images to extract the relevant measurements. To facilitate segmentation of EGF vesicles, we applied the top hat deconvolution method, to reduce the background noise and highlight bright granules. In all of the automated microscopy experiments, a minimum of 1,000 cells per biological replicate were imaged and analysed using our high-throughput microscopy set-up. No statistical methods were used to predetermine the sample sizes. The data were tested for Gaussian distribution using the Kolmogorov–Smirnov test (with a Dallal–Wilkinson–Lillie test for the *P* value). For non-Gaussian distributions, the following non-parametric tests were used: two-tailed Mann–Whitney U tests to compare two conditions or one-way analysis of variance (ANOVA; Kruskal–Wallis test) with a Dunn’s test to compare more than two data groups. For Gaussian distributions, two-tailed Student’s *t*-test were used for comparison of the means between two conditions and parametric one-way ANOVA with a Tukey’s honestly significant difference or Dunnett’s multiple comparison tests for comparisons between more than two data groups. The significance of the mean comparisons is represented on graphs by asterisks; \**P* < 0.05; \*\**P* < 0.01 and \*\*\**P* < 0.001.

### Surface biotinylation

Surface biotinylation was performed on RPE1 cells with endogenous CMG2 or on Hela cells transfected with CMG2. After 72h transfection, cells were allowed to cool down shaking at 4°C for 15 min to arrest endocytosis. Cells were then washed three times with cold PBS and treated with EZ-Link Sulfo-NHS-SS-Biotin No weight (Thermo Scientific) for 30min shaking at 4°C. Cells were then washed 3 times for 5 min with 100mM NH4Cl and lysed in IP Buffer for 1h at 4°C. Lysate were then centrifuged for 5 minutes at 5000rpm and the supernatant incubated with streptavidin agarose beads (Sigma) overnight on a wheel at 4°C. Western blots were developed using the ECL protocol (Thermo Fisher) and imaged on a Fusion Solo from Vilber Lourmat. Densitometric analysis was performed using the software Bio-1D from the manufacturer.

### Mouse experiments

ML349 was dissolved in 50%PEG300 in PBS. ML349 was injected every 2 days at 40mg/kg intraperitoneally in 11-week-old male and female C57BL/6-K18 mice ^45,46^. Control mice were injected with same volume of PEG300. After different days of treatment, different organs were collected and protein were extracted. All animal experiments were approved by Swiss cantonal veterinary office (licence VD3794b).

## Notes

### Competing Interest Statement

The authors have declared no competing interest.

## REFERENCES

1. Rahman, N. et al. The gene for juvenile hyaline fibromatosis maps to chromosome 4q21. Am J Hum Genet 71, 975–80 (2002).

2. Dowling, O. et al. Mutations in capillary morphogenesis gene-2 result in the allelic disorders juvenile hyaline fibromatosis and infantile systemic hyalinosis. Am J Hum Genet 73, 957–66 (2003).

3. Deuquet, J., Lausch, E., Superti-Furga, A. & van der Goot, F. G. The dark sides of capillary morphogenesis gene 2. Embo J 31, 3–13 (2011).

4. Burgi, J. et al. CMG2/ANTXR2 regulates extracellular collagen VI which accumulates in hyaline fibromatosis syndrome. Nature communications 8, 15861 (2017).

5. Reeves, C. V., Dufraine, J., Young, J. A. & Kitajewski, J. Anthrax toxin receptor 2 is expressed in murine and tumor vasculature and functions in endothelial proliferation and morphogenesis. Oncogene 29, 789–801 (2010).

6. Reeves, C. V. et al. Anthrax toxin receptor 2 functions in ECM homeostasis of the murine reproductive tract and promotes MMP activity. PloS one 7, e34862 (2012).

7. Schiavinato, A. et al. ANTXR2 Deficiency Promotes Cellular Senescence and Chondroid Differentiation in Hyaline Fibromatosis Syndrome Fibroblasts. J Invest Dermatol 145, 2911–2916.e7 (2025).

8. Scobie, H. M., Rainey, G. J., Bradley, K. A. & Young, J. A. Human capillary morphogenesis protein 2 functions as an anthrax toxin receptor. Proc Natl Acad Sci U S A 100, 5170–4 (2003).

9. Bradley, K. A., Mogridge, J., Mourez, M., Collier, R. J. & Young, J. A. Identification of the cellular receptor for anthrax toxin. Nature 414, 225–9. (2001).

10. Stranecky, V. et al. Mutations in ANTXR1 cause GAPO syndrome. American journal of human genetics 92, 792–9 (2013).

11. Abrami, L., Leppla, S. H. & van der Goot, F. G. Receptor palmitoylation and ubiquitination regulate anthrax toxin endocytosis. J Cell Biol 172, 309–20 (2006).

12. Anwar, M. U. et al. ER-Golgi-localized proteins TMED2 and TMED10 control the formation of plasma membrane lipid nanodomains. Developmental Cell https://doi.org/10.1016/j.devcel.2022.09.004 (2022) doi:10.1016/j.devcel.2022.09.004.

13. Mesquita, S. F. et al. Mechanisms and functions of protein S-acylation. Nat Rev Mol Cell Biol 25, 488–509 (2024).

14. Zmuda, F. & Chamberlain, L. H. Regulatory effects of post-translational modifications on zDHHC S-acyltransferases. J Biol Chem 295, 14640–14652 (2020).

15. Main, A. & Fuller, W. Protein S-Palmitoylation: advances and challenges in studying a therapeutically important lipid modification. The FEBS Journal 289, 861–882 (2022).

16. Anwar, M. U. & van der Goot, F. G. Refining S-acylation: Structure, regulation, dynamics, and therapeutic implications. J Cell Biol 222, e202307103 (2023).

17. Friebe, S., Deuquet, J. & Goot, F. G. van der. Differential Dependence on N-Glycosylation of Anthrax Toxin Receptors CMG2 and TEM8. PLOS ONE 10, e0119864 (2015).

18. Deuquet, J. et al. Systemic hyalinosis mutations in the CMG2 ectodomain leading to loss of function through retention in the endoplasmic reticulum. Hum Mutat 30, 583–9 (2009).

19. Deuquet, J. et al. Hyaline Fibromatosis Syndrome inducing mutations in the ectodomain of anthrax toxin receptor 2 can be rescued by proteasome inhibitors: Hyaline Fibromatosis Syndrome mutations. EMBO Molecular Medicine 3, 208–221 (2011).

20. Bürgi, J. et al. Ligand Binding to the Collagen VI Receptor Triggers a Talin-to-RhoA Switch that Regulates Receptor Endocytosis. Dev. Cell 53, 418–430.e4 (2020).

21. Abrami, L., Bischofberger, M., Kunz, B., Groux, R. & van der Goot, F. G. Endocytosis of the anthrax toxin is mediated by clathrin, actin and unconventional adaptors. PLoS Pathog 6, e1000792 (2010).

22. Abrami, L. et al. Palmitoylated acyl protein thioesterase APT2 deforms membranes to extract substrate acyl chains. Nat Chem Biol 17, 438–447 (2021).

23. Verkman, A. S. & Galietta, L. J. V. Chloride transport modulators as drug candidates. Am J Physiol Cell Physiol 321, C932–C946 (2021).

24. van der Sluijs, P., Hoelen, H., Schmidt, A. & Braakman, I. The Folding Pathway of ABC Transporter CFTR: Effective and Robust. J Mol Biol 436, 168591 (2024).

25. Wang, J. et al. ARF6 plays a general role in targeting palmitoylated proteins from the Golgi to the plasma membrane. J Cell Sci 136, jcs261319 (2023).

26. Abrami, L., Kunz, B. & van der Goot, F. G. Anthrax toxin triggers the activation of src-like kinases to mediate its own uptake. Proc Natl Acad Sci U S A 107, 1420–1424 (2010).

27. Castanon, I. et al. Anthrax toxin receptor 2a controls mitotic spindle positioning. Nat Cell Biol 15, 28–39 (2012).

28. Ramirez, D. M., Leppla, S. H., Schneerson, R. & Shiloach, J. Production, recovery and immunogenicity of the protective antigen from a recombinant strain of Bacillus anthracis. J Ind Microbiol Biotechnol 28, 232–238 (2002).

29. Abrami, L., Lindsay, M., Parton, R. G., Leppla, S. H. & van der Goot, F. G. Membrane insertion of anthrax protective antigen and cytoplasmic delivery of lethal factor occur at different stages of the endocytic pathway. J. Cell Biol. 166, 645–651 (2004).

30. Won, S. J. et al. Molecular Mechanism for Isoform-Selective Inhibition of Acyl Protein Thioesterases 1 and 2 (APT1 and APT2). ACS chemical biology 11, 3374–3382 (2016).

31. Moayeri, M., Leppla, S. H., Vrentas, C., Pomerantsev, A. P. & Liu, S. Anthrax Pathogenesis. Annu Rev Microbiol 69, 185–208 (2015).

32. Friebe, S., van der Goot, F. G. & Bürgi, J. The Ins and Outs of Anthrax Toxin. Toxins (Basel) 8, (2016).

33. Abrami, L. et al. Hijacking Multivesicular Bodies Enables Long-Term and Exosome-Mediated Long-Distance Action of Anthrax Toxin. Cell reports 27, 986–996 (2013).

34. Bürgi, J., Xue, B., Uversky, V. N. & van der Goot, F. G. Intrinsic Disorder in Transmembrane Proteins: Roles in Signaling and Topology Prediction. PLoS ONE 11, e0158594 (2016).

35. Blanc, M. et al. SwissPalm: Protein Palmitoylation database. F1000Research 4, 261 (2015).

36. Blanc, M., David, F. P. A. & van der Goot, F. G. SwissPalm 2: Protein S-Palmitoylation Database. Methods Mol. Biol. 2009, 203–214 (2019).

37. Blaskovic, S., Blanc, M. & van der Goot, F. G. What does S-palmitoylation do to membrane proteins? The FEBS journal 280, 2766–2774 (2013).

38. Fukata, Y., Iwanaga, T. & Fukata, M. Systematic screening for palmitoyl transferase activity of the DHHC protein family in mammalian cells. Methods 40, 177–82 (2006).

39. Deuquet, J. et al. Hyaline Fibromatosis Syndrome inducing mutations in the ectodomain of anthrax toxin receptor 2 can be rescued by proteasome inhibitors. EMBO Mol Med 3, 208–221 (2011).

40. Feld, G. K., Kintzer, A. F., Tang, I., Thoren, K. L. & Krantz, B. A. Domain flexibility modulates the heterogeneous assembly mechanism of anthrax toxin protective antigen. Journal of molecular biology 415, 159–74 (2012).

41. Colombatti, A., Mucignat, M. T. & Bonaldo, P. Secretion and matrix assembly of recombinant type VI collagen. J Biol Chem 270, 13105–13111 (1995).

42. Forrester, M. T. et al. Site-specific analysis of protein S-acylation by resin-assisted capture. J Lipid Res 52, 393–8 (2011).

43. Abrami, L., Denhardt-Eriksson, R. A., Hatzimanikatis, V. & van der Goot, F. G. Dynamic Radiolabeling of S-Palmitoylated Proteins. Methods Mol. Biol. 2009, 111–127 (2019).

44. Brankatschk, B. et al. Regulation of the EGF transcriptional response by endocytic sorting. Sci Signal 5, ra21 (2012).

45. Sun, S.-H. et al. A Mouse Model of SARS-CoV-2 Infection and Pathogenesis. Cell Host Microbe 28, 124–133.e4 (2020).

46. Mesquita, S. F. et al. SARS-CoV-2 hijacks a cell damage response, which induces transcription of a more efficient Spike S-acyltransferase. Nat Commun 14, 7302 (2023).

